# Nanometer-Scale Imaging of Compartment-Specific Localization and Dynamics of Voltage-Gated Sodium Channels

**DOI:** 10.1101/2021.09.28.462114

**Authors:** Hui Liu, Hong-Gang Wang, Geoffrey S. Pitt, Zhe J. Liu

## Abstract

Membrane excitability and cell-to-cell communication in the brain are tightly regulated by diverse ion channels and receptor proteins localized to distinct membrane compartments. Currently, a major technical barrier in cellular neuroscience is lack of reliable methods to label these membrane proteins and image their sub-cellular localization and dynamics. To overcome this challenge, we devised optical imaging strategies that enable systematic characterization of subcellular composition, relative abundances and trafficking dynamics of membrane proteins at nanometer scales in cultured neurons as well as in the brain. Using these methods, we revealed exquisite developmental regulation of subcellular distributions of voltage-gated sodium channel (VGSC) Na_v_1.2 and Na_v_1.6, settling a decade long debate regarding the molecular identity of sodium channels in dendrites. In addition, we discovered a previously uncharacterized trafficking pathway that targets Na_v_1.2 to unmyelinated fragments in the distal axon. Myelination counteracts this pathway, facilitating the installment of Nav1.6 as the dominant VGSC in the axon. Together, these imaging approaches unveiled compartment-specific trafficking mechanisms underpinning differential membrane distributions of VGSCs and open avenues to decipher how membrane protein localization and dynamics contribute to neural computation in the brain.

## INTRODUCTION

Information processing in the brain is regulated at the molecular level by diverse membrane proteins such as ion channels and receptor proteins (Catterall, Goldin, & Waxman, 2005; Hodgkin & Huxley, 1952). Two broad types of ion channels are: voltage-gated ion channels and ligand-gated ion channels. In mammals, it was estimated that ∼140 genes encode voltage-gated K^+^, Na^+^ and Ca^2+^ channels (Yu, Yarov-Yarovoy, Gutman, & Catterall, 2005). In addition to ion channels, a large number of metabotropic receptors convert extracellular stimuli to intracellular signaling responses via G protein or protein kinase mediated pathways (Niswender & Conn, 2010; Stevens et al., 2013). Single-cell RNA-seq data reveal that neuron and glial populations in the brain express distinct sets of membrane proteins (Cembrowski, Wang, Sugino, Shields, & Spruston, 2016; Zeisel et al., 2015), suggesting that membrane physiology is tightly regulated in a cell-type specific manner. Thus, one key challenge in cellular neuroscience is to decipher how the spatial distribution of ion channels and receptor proteins along the complex membrane topology controls signal integration, action potential initiation, backward propagation and synaptic plasticity in the context of a specific circuitry (Lai & Jan, 2006; Vacher, Mohapatra, & Trimmer, 2008).

Currently, one of the major challenges to probe membrane proteins in the brain is lack of reliable methods to label endogenous ion channel or receptor proteins. Specifically, traditional immuno-labeling is associated with several limitations: 1) nonspecific cross-reaction, especially for antibodies against closely related channels and receptors; 2) insufficient sensitivity when the copy number of the target protein is low; 3) subcellular localization information is obscured by high packing density of neurites in the brain (Mikuni, Nishiyama, Sun, Kamasawa, & Yasuda, 2016). As a result, mapping sub-cellular localization of membrane proteins poses tremendous challenges for neuroscientists (Baker, 2020).

To address these limitations, here we combined CRISPR/Cas9 *in vivo* genome editing with high affinity peptide tags (V5 (GKPIPNPLLGLDST) or HA (YPYDVPDYA)) and self-labeling tags (*e.g.* HaloTag) to label membrane proteins. Sparse cell labeling and high sensitivity of monoclonal antibodies enable us to reconstruct subcellular localizations of membrane proteins with high spatial resolution. Using brain-enriched voltage-gated sodium channel Na_v_1.2 and Na_v_1.6 as the model, we found that Na_v_1.2 is highly enriched in the AIS, dendrites and unmyelinated distal axon branches during early development. As animals develop into adults, Na_v_1.6 levels increase while Na_v_1.2 levels decrease in dendrites, accompanied by myelination dependent exclusion of Na_v_1.2 from the axon and an eventual installment of Na_v_1.6 as the dominant VGSC at the AIS and nodes of Ranvier. Super resolution and live-cell single molecule imaging in cultured neurons enables real time investigation of VGSC trafficking dynamics at nanometer scales. We found that while localization of Na_v_1.2 and Na_v_1.6 to the AIS is dependent on Ankyrin G binding domain (ABD) as previously described (Garrido et al., 2003; Lemaillet, Walker, & Lambert, 2003), the targeting of Na_v_1.2 to unmyelinated fragments in the distal axon requires separated signals within the intracellular loop 1 (ICL1) between transmembrane domain I and II. Specifically, Na_v_1.2 ICL1 suppresses AIS retention and permits the membrane loading of Na_v_1.2 at the distal axon. Together, these results unveiled compartment-specific localization and trafficking mechanisms for Na_v_1.2 and Na_v_1.6, which could be modulated independently to fine tune membrane composition and physiological functions of VGSCs in the brain.

## RESULTS

### Differential subcellular localizations of Na_v_1.2 and Na_v_1.6 in cultured hippocampal neurons

Brain enriched VGSC Na_v_1.2 and Na_v_1.6 are critical for electrical signaling in the central nervous system and their mutations are associated with human genetic diseases such as infant epilepsy and autism spectrum disorder (Meisler, Hill, & Yu, 2021; Sanders et al., 2018). Previous studies indicated the prominent presence of Na_v_1.2 and Na_v_1.6 in the axon initial segment (AIS) of excitatory neurons. However, their composition in other neuronal compartments remains unclear (Johnson, Herold, Milner, Hemmings, & Platholi, 2017; Lorincz & Nusser, 2010; Spratt et al., 2019). Specifically, because of relative low copy number of VGSCs in dendrites and insufficient sensitivity of traditional methods, the dendritic composition of VGSCs is still under debate. For example, Na_v_1.6 was shown to localize in dendrites of hippocampal CA1 pyramidal neurons via a highly sensitive electron microscopic immune-gold technique, by which Na_v_1.2 was not detected (Lorincz & Nusser, 2010). However, another study showed the presence of Na_v_1.2 in dendrites and spines in the hippocampal CA1 region (Johnson et al., 2017). In addition, electrophysiology studies indicated that Na_v_1.2 plays a key role in Na_v_ currents at the somatodendritic region of cortical neurons (Hu et al., 2009; Spratt et al., 2019).

To resolve this debate, we took advantage of previously established homology-independent targeted integration (HITI) genome editing method (Suzuki et al., 2016) (**Figure 1-figure supplement 1A**) and tagged *Scn2a* (Na_v_1.2) and *Scn8a* (Na_v_1.6) with small peptide tags (V5 or HA). We reason that the small size of these tags would minimize the risk of perturbing their physiological functions. Indeed, V5 tag insertion at two independent locations (C-terminus versus the extracellular loop between segment 5 and 6 in domain I) gave rise to comparable subcellular localization patterns (**Figure 1-figure supplement 2 and 3**). Single cell recording confirmed that the tag insertion did not affect electrophysiological properties of Na_v_1.2 and Na_v_1.6 (**Figure 1- figure supplement 5**). Consistent with previous immune-staining results (Xu, Zhong, & Zhuang, 2013), super-resolution STED imaging revealed that tagged Na_v_1.2 and Na_v_1.6 form ∼200 nm periodic striations that showed anti-phased exclusion from actin rings at the AIS, further validating the labeling strategy (**Figure 1B, C**).

**Figure 1.**
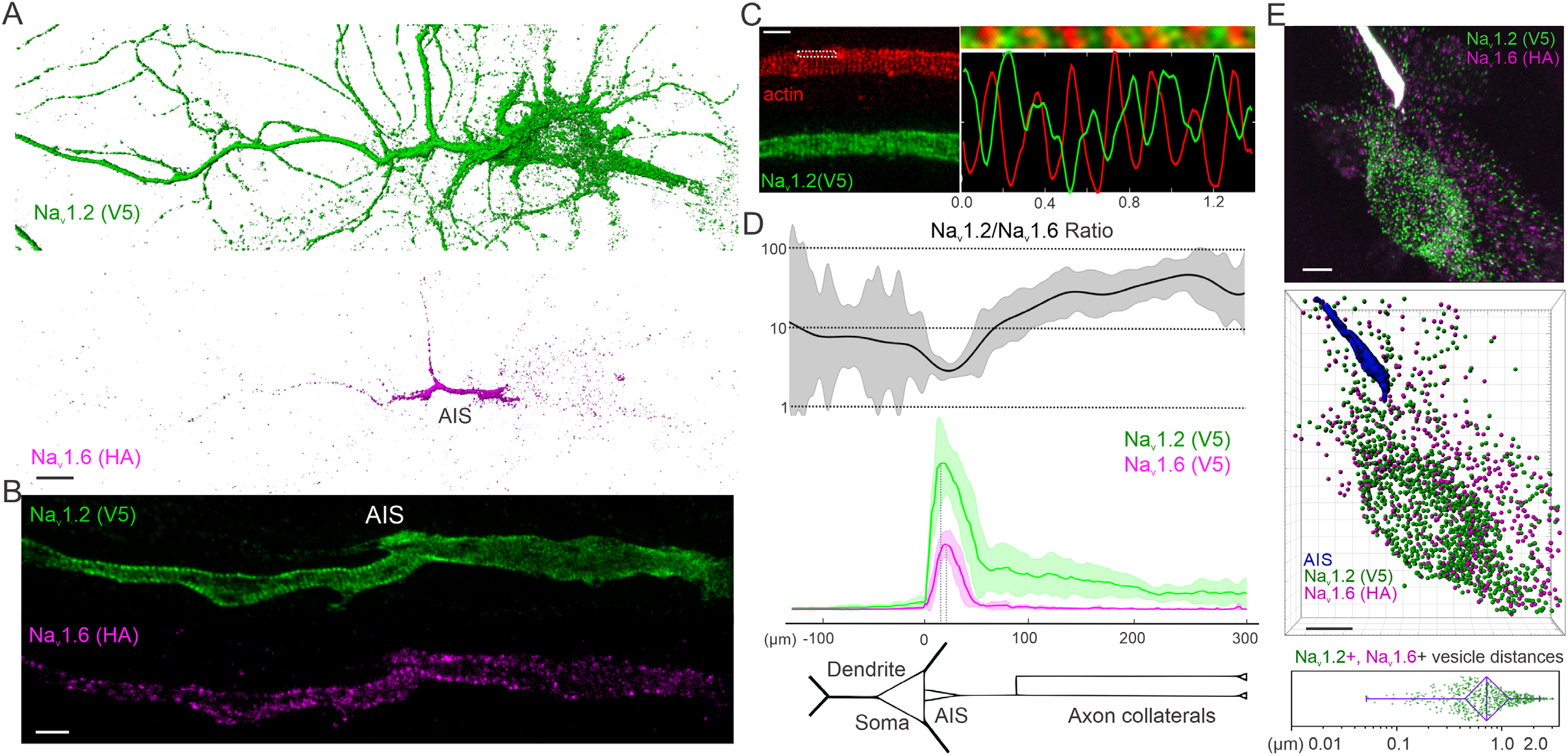
Sub-cellular distributions of Na_v_1.2 and Na_v_1.6 in cultured hippocampal neurons. (A) Labeling of Na_v_1.2 with V5 tag and Na_v_1.6 with HA tag in the same neuron. Scale bar, 10 µm. (B) Super-resolution STED imaging of labeled Na_v_1.2 and Na_v_1.6 in the AIS region. Scale bar, 1 µm. (**C**) Anti-phase periodic striations of V5-labeled Na_v_1.2 and actin in the AIS region. The top image on the right shows a zoom-in view of the rectangle region with dashed lines in the left image. Bottom shows the intensity curves of Na_v_1.2 (green) and actin (red) along the horizontal line. Scale bar, 1 µm. (**D**) Analysis of Na_v_1.2 (n = 12) and Na_v_1.6 (n = 14) relative intensities along the dendrite and axon of cultured hippocampal neurons (Figure 1-source data 1). The top curve shows the Na_v_1.2/Na_v_1.6 intensity ratio calculated by using Na_v_1.2 and Na_v_1.6 intensity data shown in the middle panel. Error bars (shadow areas) represent SD. (**E**) Airyscan imaging of Na_v_1.2- and Na_v_1.6-positive vesicles in the soma. The middle panel shows computer aided segmentation of Na_v_1.2 and Na_v_1.6 vesicles and the bottom box chart shows the distribution of physical distances between these two vesicle populations (Figure 1-source data 2). Scale bar, 5 µm. In the box chart, right and left error bars represent 95% and 5% percentile, respectively; triangle represents the range from 25% to 75% percentile; center line represents the median.

To quantify relative abundance of Na_v_1.2 and Na_v_1.6 in distinct neuronal compartments, we tagged both channels with the V5 tag, followed by labeling and imaging under the same condition. By cross referencing with AIS (Ankyrin G) and dendrite (MAP2) markers, we found that both channels showed highest enrichment in the AIS (**Figure 1-figure supplement 3**), consistent with previous reports (Hu et al., 2009; Lorincz & Nusser, 2010). Interestingly however, we found that the relative abundance of Na_v_1.2 are much higher than Na_v_1.6 in the distal axon and dendrites (**Figure 1D, Figure 1-figure supplement 4**). To confirm that what observed are not influenced by cell-type specific expression, we employed sequential HITI editing and achieved dual labeling of Na_v_1.2 (V5) and Na_v_1.6 (HA) in the same cell population (**Figure 1-figure supplement 1B**). Na_v_1.2 and Na_v_1.6 staining patterns in the dual labeling condition were consistent with what were observed in separate populations with highest levels of Na_v_1.2 and Na_v_1.6 in the AIS and Na_v_1.2 as the dominant VGSC in the distal axon and dendrites (**Figure 1A, Figure 1-figure supplement 1B, C, Video 1**). Live-cell non-permeable staining confirmed that Na_v_1.2 is indeed inserted into cell membrane in the distal axon and dendrites (**Figure 1-figure supplement 2**). Thus, here we were able to unambiguously confirm the localization of Na_v_1.2 in dendrites of cultured hippocampal pyramidal neurons.

The differential localization patterns of Na_v_1.2 and Na_v_1.6 prompted us to probe underlying trafficking mechanisms. Super-resolution Airyscan imaging and computer-aid segmentation revealed no co-labelled fraction between Na_v_1.2 and Na_v_1.6 positive trafficking vesicles (**Figure 1E, Video 2**), suggesting that once synthesized, Na_v_1.2 and Na_v_1.6 are sorted into distinct vesicle populations potentially coupled with separated trafficking and membrane loading pathways.

### Developmental regulation of Na_v_1.2 and Na_v_1.6 subcellular localizations in vivo

To map Na_v_1.2 and Na_v_1.6 localizations *in vivo*, we used *in utero* electroporation to deliver the HITI construct into heterozygous H11-SpCas9 mouse embryos expressing Cas9 in all cell types (Chiou et al., 2015). The resulting sparse neuron labeling in the mouse cortex and hippocampus enabled us to quantify their levels in individual neurites across large distances at different developmental stages (Postnatal (P) 15, P30 (∼one month) and P90 (∼three months)) (**Figure 2A, Video 3-5**). In agreement with previous reports (Hu et al., 2009; Tian, Wang, Ke, Guo, & Shu, 2014; Yamagata, Ogiwara, Mazaki, Yanagawa, & Yamakawa, 2017), we found that Na_v_1.2 and Na_v_1.6 are mainly expressed in CaMKIIα-positive excitatory neurons, with no detectable expression in GAD67-positive inhibitory neurons (**Figure 2-figure supplement 2C-E**).

**Figure 2.**
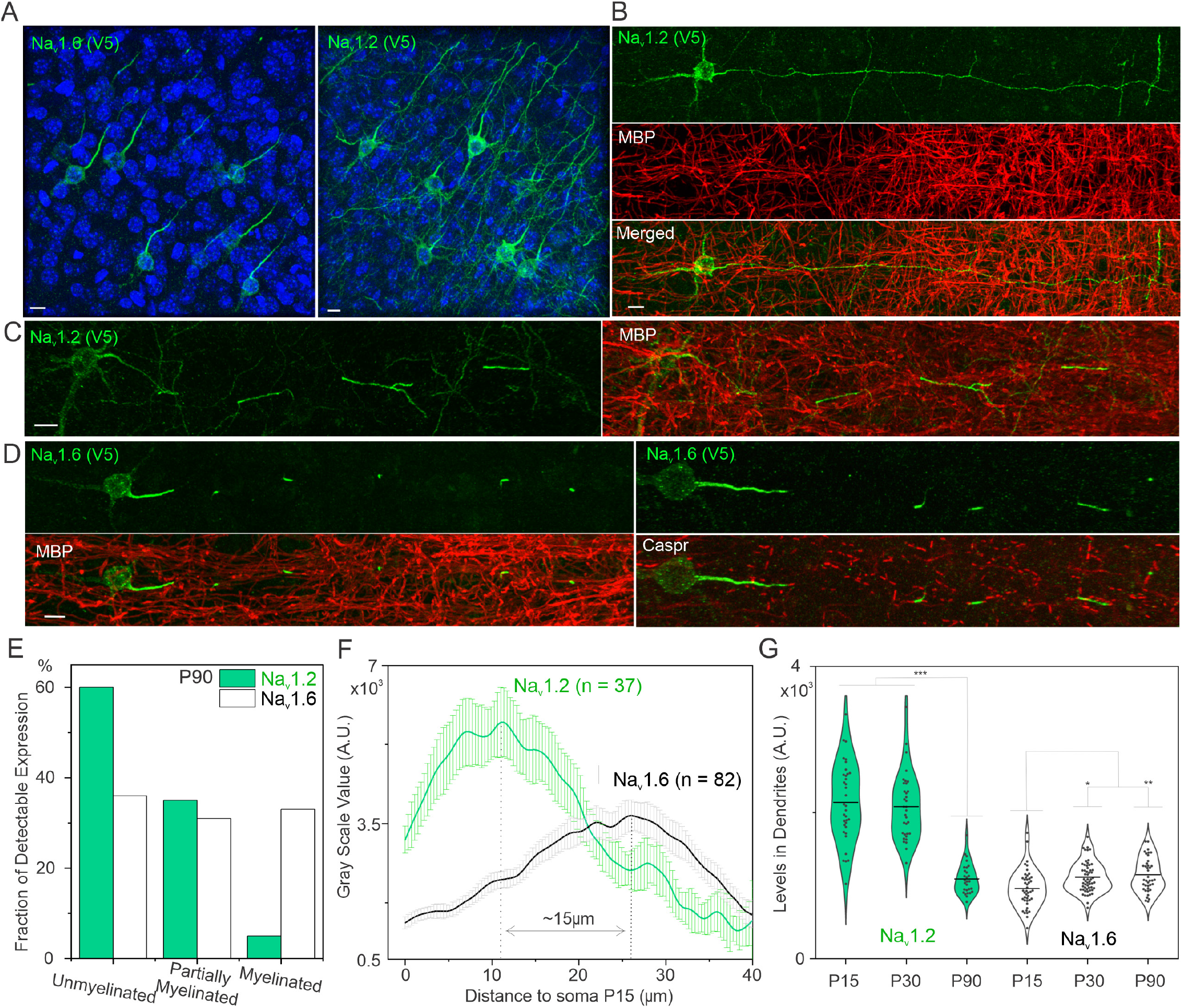
Developmental regulation of Na_v_1.2 and Na_v_1.6 localization and expression in the brain. (**A**) Representative images of Na_v_1.2 and Na_v_1.6 labeled with V5 tag in the cortex. Blue channel shows the Hoechst stain. Scale bar, 10 µm. (**B** and **C**) Double-immunostaining of V5-labeled Na_v_1.2 with MBP (red) in unmyelinated neuron (**B**) and partially myelinated neuron (**C**). Scale bar, 10 µm. (**D**) Double-immunostaining of V5-labeled Na_v_1.6 with MBP (left) and Caspr (right). Scale bar, 10 µm. (**E**) The percentage of unmyelinated, partially myelinated, and myelinated neurons in Na_v_1.2- or Na_v_1.6-positive knockin neurons. The number of cells analyzed: Na_v_1.2, 60; Na_v_1.6, 97. (**F**) Intensity measurements of V5-labeled Na_v_1.2 and Na_v_1.6 in the AIS of cortical neurons at P15 (Figure 2-source data 1). Error bars represent SEM; n indicates the number of cells analyzed. (**G**) Violin plots of intensity measurements of V5-labeled Na_v_1.2 and Na_v_1.6 in dendrites of cortical neurons at different ages (Figure 2-source data 2). In the graph, the line represents the mean. *, *p*-value < 0.05; **, *p*-value < 0.01; ***, *p*-value < 0.001.

Because both Na_v_1.2 and Na_v_1.6 were tagged with the V5 peptide, we were able to estimate their relative abundances with high spatial resolution. At P15, both Na_v_1.2 and Na_v_1.6 were enriched at the AIS (**Figure 2A, Figure 2-figure supplement 2A, B**) with higher levels of Na_v_1.2 in the distal axon and dendrites (**Figure 2A, G**), similar to our observations in cultured hippocampal neurons. Interestingly, Na_v_1.2 is enriched at the proximal part of the AIS while Na_v_1.6 is concentrated at the distal part of the AIS with a ∼15 µm gap between their concentration peaks (**Figure 2F**), consistent with a previous report (Hu et al., 2009). Interestingly however, Na_v_1.2 levels decreased significantly at the proximal AIS with the Na_v_1.6 concentration peak shifting inwards at P30 and P90 (**Figure 2-figure supplement 1**).

We were also able to confirm the localization of Na_v_1.2 in dendrites of cortical and hippocampal pyramidal neurons *in vivo* (**Figure 2A, Figure 2-figure supplement 2A**). In addition, we found that Na_v_1.2 is the dominant VGSC in dendrites during early development and its concentration gradually decreases, accompanied by an increase of Na_v_1.6 levels at this region as mice mature (**Figure 2G**).

Previous electrophysiology experiments revealed that Na_v_1.6 has much lower activation threshold and larger persistent currents than Na_v_1.2 (Rush, Dib-Hajj, & Waxman, 2005). These results suggested that neurons actively adjust their excitability by fine tuning the membrane composition and localization of Na_v_1.2 and Na_v_1.6 at different developmental stages. Specifically, during early development, the information processing at the somatodendritic region and the proximal AIS is mainly mediated by Na_v_1.2, while Na_v_1.6 plays increasing important roles at these regions as the animal matures. Consistent with these results, human genetic studies found that mutations in Na_v_1.2 are primarily associated with early developmental diseases such as infant epilepsy and autism spectrum disorder (Meisler et al., 2021; Sanders et al., 2018).

### Myelination status as a key indicator of Na_v_1.2 and Na_v_1.6 localization patterns

One consistent observation across all developmental stages is that the axonal coverage by Na_v_1.2 is largely uninterrupted in neurons with high Na_v_1.2 expression levels, suggesting that these cells are unmyelinated (**Figure 2A, B, Video 3**). Indeed, when using myelin basic protein (MBP) to co-stain samples, we found that Na_v_1.2 was preferentially expressed (∼60%) in unmyelinated neurons with a smaller fraction (∼35%) of detectable expression in partially myelinated neurons and the lowest fraction (∼5%) in fully myelinated neurons (**Figure 2B, C, E**). By contrast, Na_v_1.6 has similar fractions of detectable expression across all 3 populations (**Figure 2E**). In addition, we found that the axonal coverage by Na_v_1.6 is restricted to Ankyrin G positive regions such as the AIS and nodes of Ranvier (**Figure 2D, Video 4 and Video 5**), whereas Na_v_1.2 broadly covers unmyelinated axonal fragments (**Figure 2B, Video 3**). Our results support that localizations and expression levels of these two channels alter with the myelination status of a neuron. Specifically, myelination excludes Na_v_1.2 and decreases its expression levels in the axon, with an eventual installment of Na_v_1.6 as the dominant VGSC at the AIS and nodes of Ranvier in fully myelinated neurons. Previous reports showed that, upon neuronal injury, the large persistent currents of Na_v_1.6 at demyelinated sites trigger reverse action of Na^+^-Ca^2+^ exchanger, leading to Ca^2+^ influx that further damages the axon (Craner et al., 2004; Rush et al., 2005). This result explains the physiological need of coating unmyelinated axonal fragments with a VGSC that conducts smaller persistent currents such as Na_v_1.2.

### Compartment-specific Targeting Mechanisms for Na_v_1.2 and Na_v_1.6

The differential localization patterns and separated vesicle populations associated with Na_v_1.2 and Na_v_1.6 suggest that they are trafficked by different pathways (**Figure 1E and 2A**). To study the underlying mechanism, we established a 2-color imaging assay in which Na_v_1.2 and Na_v_1.6 (tagged with V5 and HA respectively) were co-expressed in cultured hippocampal neurons where their localization patterns can be directly compared in distinct subcellular compartments. We found that exogenously expressed Na_v_1.2(V5) and Na_v_1.6(HA) displayed similar localization patterns as endogenous knock in proteins, with both VGSCs enriched in AIS and Na_v_1.2 as the dominant VGSC in distal axon and dendrites (**Figure 3-figure supplement 1A**). Because of conserved sequence homology in membrane embedded domains, we focused on intracellular loops which have larger sequence divergences. Consistent with previous reports (Garrido et al., 2003; Gasser et al., 2012; Lemaillet et al., 2003), we found that ABD deletion led to the loss of Na_v_1.2 and Na_v_1.6’s enrichment at the AIS, interestingly with no significant effects on their distal axon and dendrite localization patterns (**Figure 3A, B, Figure 3-figure supplement 1B**). This result suggests that localization of Na_v_1.2 to the distal axon and dendrites is independent of its AIS anchoring signal (ABD). By extensive domain swapping between Na_v_1.2 and Na_v_1.6 (**Figure 3-figure supplement 2**), we identified the intracellular loop 1 (ICL1) between transmembrane domain I and II (**Figure 3A, Figure 3-figure supplement 4A**) as a key determinant for selective enrichment of Na_v_1.2 in the distal axon. Specifically, Na_v_1.2 with Na_v_1.6 ICL1 displayed the same localization pattern as Na_v_1.6, with very low enrichment in the distal axon. Conversely, Na_v_1.6 harboring ICL1 from Na_v_1.2 showed comparable enrichment in the distal axon as Na_v_1.2 (**Figure 3C, Figure 3-figure supplement 1A and 2C**). These results suggest that Na_v_1.2 ICL1 contains previously uncharacterized distal axon targeting and membrane loading signals.

**Figure 3.**
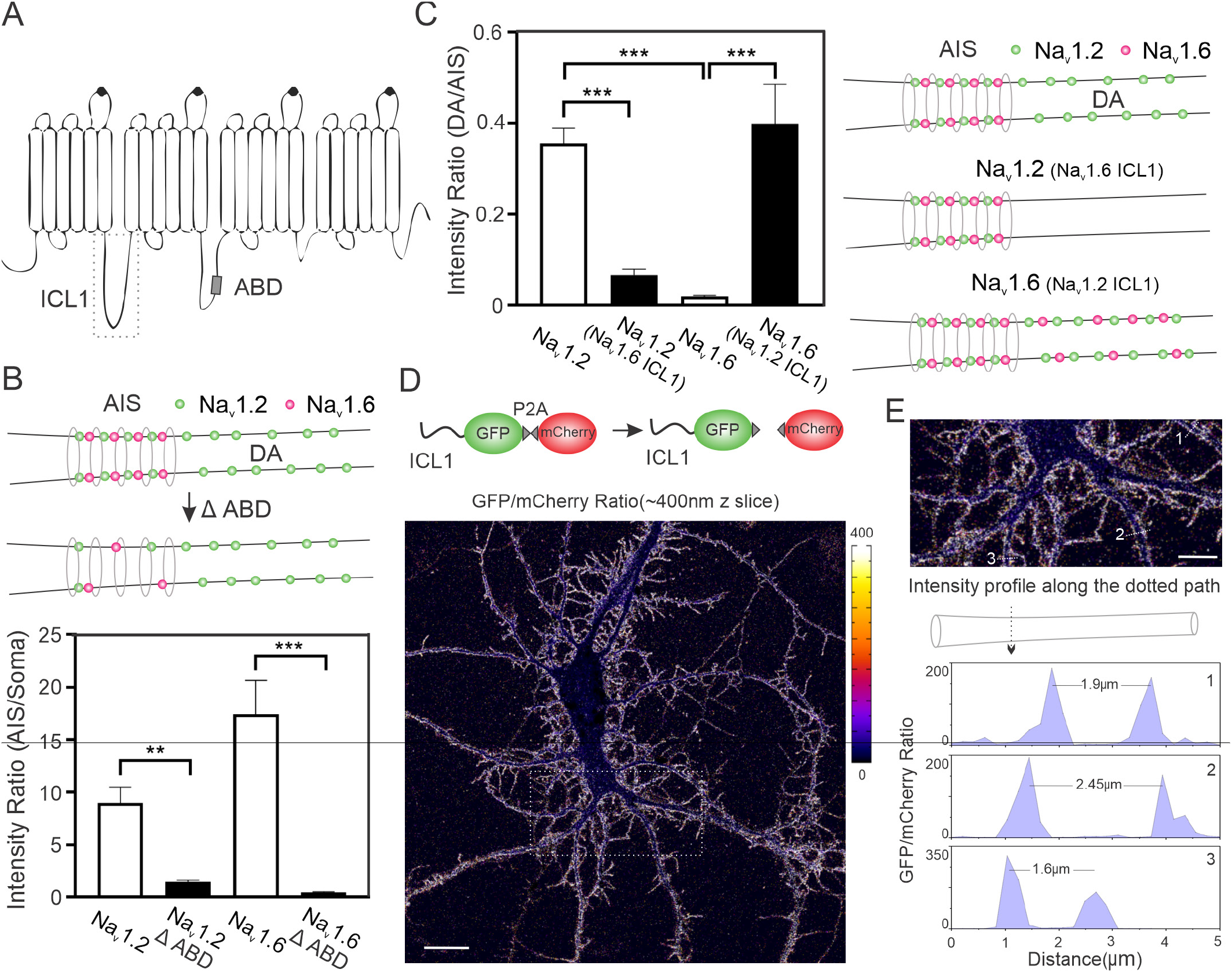
Compartment-specific targeting mechanisms for Na_v_1.2 and Na_v_1.6. (**A**) Rectangle region with dashed lines shows the ICL1 region of Na_v_1.2 and Na_v_1.6. ABD is indicated by a small gray rectangle. (**B**) Deletion of ABD abolished enrichment of Na_v_1.2 and Na_v_1.6 in AIS. Top shows the cartoon demonstration and bottom shows the statistical analysis of the intensity ratio between AIS and soma after ABD deletion (Figure 3-source data 1). (**C**) Na_v_1.2 with Na_v_1.6 ICL1 showed dramatic less enrichment in the distal axon. Conversely, Na_v_1.6 with Na_v_1.2 ICL1 gains the ability to localize to the distal axon. Left shows the statistical analysis of the intensity ratio between distal axon and soma of Na_v_1.2 or Na_v_1.6 after ICL1 replacement (Figure 3-source data 2) and right shows the cartoon demonstration. Error bars represent SEM. **, *p*-value < 0.01; ***, *p*-value < 0.001. (**D**) Top shows the illustration of the ratiometric localization analysis. After translation, ICL1-GFP and mCherry proteins are separated due to ribosome skipping at P2A. A representative intensity ratio (Na_v_1.2 ICL1-GFP/mCherry) image showed the localization of Na_v_1.2 ICL1 to membrane. Right color bar indicates the ratio level. Scale bar, 20 µm. (**E**) Top shows the enlarged view of the rectangle region with dashed lines in (**D**). Three neurites were chosen to analyze their intensity profiles along the dotted paths, of which the results were shown below. Scale bar, 10 µm.

To further dissect the function of ICL1, we fused it to GFP-P2A-mCherry. Strikingly, we found that Na_v_1.2 ICL1 itself was able to broadly target GFP to cell membrane across different compartments (soma, axon and dendrites) (**Figure 3D, E, Figure 3-figure supplement 3A**), whereas Na_v_1.6 ICL1-GFP signals were largely in the nucleus (**Figure 3-figure supplement 3B-D**), consistent with previous reporting of a nucleus localization signal within this region (Onwuli et al., 2017). Using this assay, we further determined that a 36 amino acid region (AA725-760) within Na_v_1.2 ICL1 was sufficient for anchoring GFP to membrane (**Figure 3-figure supplement 4B**). Together, these results suggest that Na_v_1.2 is targeted to AIS and the distal axon via a separated, previously characterized trafficking pathway.

### A model for targeting Na_v_1.2 to unmyelinated fragments in the distal axon

To build a physical model to explain how differential subcellular localizations of Na_v_1.2 and Na_v_1.6 are dynamically established at the molecular level, we sought to utilize live-cell single-molecule imaging approaches that we established previously (Chen et al., 2014; Liu et al., 2018). These methods with nanometer scale detection sensitivity have been widely adopted to study transcription factor and vesicle dynamics (Chen et al., 2014; Chong et al., 2018; Knight et al., 2015; Liu et al., 2018) in live cells. To achieve live-cell labeling, we knocked in HaloTag at the C-terminus of these two VGSCs, followed by staining with bright, membrane permeable Janelia Fluor dyes (Grimm et al., 2015). We found that localization patterns of HaloTag-labeled Na_v_1.2 and Na_v_1.6 in the AIS were similar to these tagged with V5 and HA tag (**Figure 4-figure supplement 1**), suggesting that HaloTag labeling did not significantly perturb Na_v_1.2 and Na_v_1.6 trafficking. Then, we devised a pulse-chase assay in which we first used high concentrations of JF646-HaloTag ligand (HTL) to block pre-existing VGSC-HaloTag molecules and then we pulsed cells with JF549-HTL for short durations to label newly synthesized VGSCs. This technique allowed us to control labeling density by tuning pulse durations and thus obtain long trajectories of trafficking VGSCs under sparse labeling conditions (**Figure 4A**).

**Figure 4.**
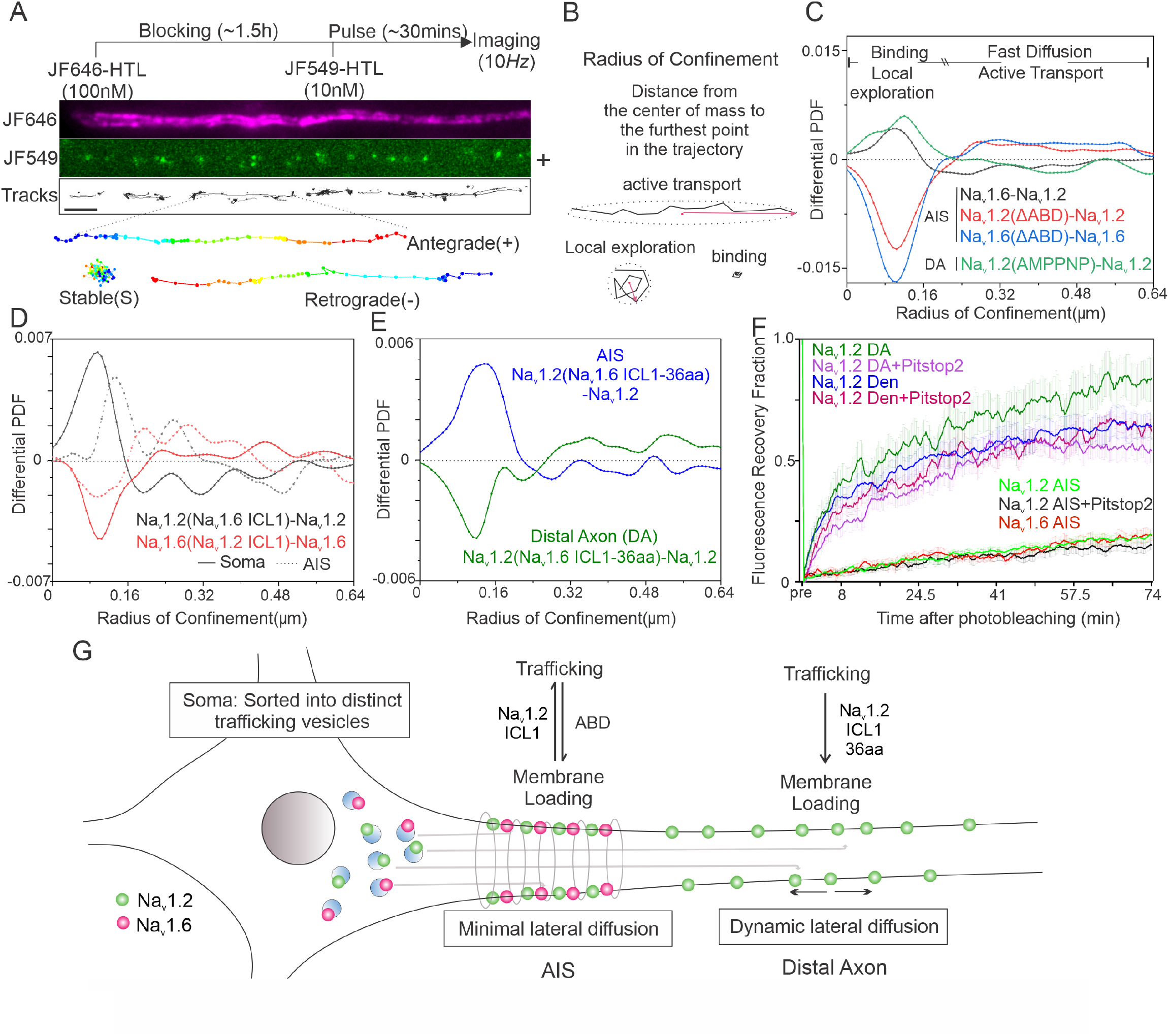
Live Imaging of Na_v_1.2 and Na_v_1.6 trafficking and lateral diffusion dynamics. (**A**) Pulse-chase single molecule imaging of Na_v_1.2 and Na_v_1.6. Top shows the experimental flowchart. Middle shows a representative AIS image, with JF646 bulk labeling image, JF549 pulse-chased single molecule signals and analyzed single molecule moving trajectories. Bottom shows three different types of trajectories: stable binding, anterograde and retrograde movement (blue to red color change represents time progression). (**B**) Definition of Radius of Confinement (RC) for analyzing single molecule moving dynamics. Stable binding and local exploration events have smaller RCs, whereas active transport and fast diffusion events should have larger RCs. (**C**) Comparative RC distribution curves of Na_v_1.6 - Na_v_1.2 (black curve), Na_v_1.2(ΔABD) - Na_v_1.2 (red curve), Na_v_1.6(ΔABD) - Na_v_1.6 (blue curve) in the AIS region and Na_v_1.2(AMPPNP) - Na_v_1.2 (green curve) in the distal axon (Figure 4-source data 1). Differential PDF = 0 stands for equal fraction. (**D**) Comparative RC distribution curves of Na_v_1.2(Na_v_1.6 ICL1) – Na_v_1.2 (black curve) and Na_v_1.6(Na_v_1.2 ICL1) – Na_v_1.6 (red curve) in soma (solid line) and AIS (dotted line) (Figure 4-source data 2). (**E**) Comparative RC distribution curves of Na_v_1.2(Na_v_1.6 ICL1-36aa) – Na_v_1.2 in AIS (blue) and distal axon (green) (Figure 4-source data 3). (**F**) Fluorescent recovery curves of Na_v_1.2 and Na_v_1.6 in different neuronal compartments after photobleaching (Figure 4-source data 4). Pitstop 2 (30 µM) was added to inhibit endocytosis during the labeling and imaging period. Error bars represent SEM. (**G**) A cartoon model showing that Na_v_1.2 ICL1 is important for suppressing AIS anchoring and facilitating membrane insertion at the distal axon.

To establish a simple and effective method to quantify dynamic states (stable binding, local exploration, diffusion and active transport) associated with trafficking and membrane loading, we took advantage of the “Radius of Confinement (RC)” parameter which we used successfully to study binding and diffusion states of diverse transcription factors (Lerner et al., 2020). Specifically, the RC is defined as the distance from the center of mass to the furthest point of the trajectory (**Figure 4B**). Intuitively, fast diffusion and active transport events along the neurites should correlate with larger RCs compared with bound and local exploration states (**Figure 4B, C**). Indeed, we found that ABD deletion in Na_v_1.2 or Na_v_1.6 led to a dramatic reduction of shorter RC fractions and an increase in longer RC fractions, reflecting less binding but more active transport events in the AIS, consistent with known functions of ABD (**Figure 4C**). Conversely, inhibiting active transport by ATP analog (AMPPNP) significantly reduced active transport (longer RC) fractions but increased short RC fractions in distal axon (**Figure 4C**), confirming the ability of the RC analysis to separate distinct dynamic states.

To dissect the molecular basis underlying each dynamic state, we next coupled the RC analysis with genetic perturbations. We found that Na_v_1.2 displayed significantly less binding and more active transport events in AIS than Na_v_1.6 (**Figure 4C**). Similarly, replacing ICL1 in Na_v_1.6 with Na_v_1.2 ICL1 deceased binding and induced more active transport in the AIS and soma. The opposite is true as Na_v_1.2 with Na_v_1.6 ICL1 has more binding but less active transport events than Na_v_1.2 (**Figure 4D**). The remarkable consistency in these results support that Na_v_1.2 ICL1 promotes active transport and suppresses retention in the AIS, counterbalancing the anchoring effect of ABD. Complementary with these results, we found that Na_v_1.2 with the membrane anchoring domain ICL-36aa (AA725-760) replaced with the same region from Na_v_1.6 showed much less binding at the distal axon, suggesting that this domain is critical for membrane insertion of Na_v_1.2 (**Figure 4E**), consistent with its ability to anchor GFP to cell membrane. Taken together, these results suggest that localization of Na_v_1.2 to the distal axon requires two distinct functions of ICL1: one for reducing anchoring at the AIS; the other for promoting membrane insertion at the distal axon (**Figure 4G**).

Next, we used fluorescence recovery after photobleaching (FRAP) to examine lateral diffusion of membrane bound Na_v_1.2 and Na_v_1.6 across different compartments. For this assay, we utilized Pitstop 2 to block endocytosis mediated exchanges on the membrane so that fluorescent recovery is largely dependent on lateral diffusion. We found that both Na_v_1.2 and Na_v_1.6 showed slow exchanging rates at the AIS with only ∼15% recovery 1 hour after photobleaching, whereas, in the distal axon and dendrites, VGSC is more dynamic, with ∼60 percent recovery in the distal axon and ∼50 percent recovery in the dendrites ∼0.5 hour after photobleaching (**Figure 4F, Video 6**). These results are consistent with that lateral diffusion of VGSCs is also regulated in a compartment-specific fashion ranging from minimal mobility in the AIS to faster diffusion in dendrites and the distal axon (**Figure 4G**). It is likely that the reduced VGSC lateral diffusion in the AIS could be related to the unique, ring-like cytoskeleton structures at this region showing anti-phase, exclusive distributions to VGSC striations (**Figure 1B, C**).

## DISCUSSION

Here, we demonstrated nanometer-scale imaging strategies to characterize sub-cellular localization, relative abundances and tracking dynamics of membrane proteins in the brain. We overcame the limitations of traditional immune-labeling methods and provided an unprecedentedly clear view of Na_v_1.2 and Na_v_1.6 subcellular localizations both *in vitro* and *in vivo*. Our results confirmed key results from previous studies and provided new insights into compartment-specific VGSC localization patterns at different developmental stages, providing direct imaging evidence to clarify decade long debates in the field.

The most pronounced difference between Na_v_1.2 and Na_v_1.6 localizations that we observed is in the distal axon, where their expression and localization patterns showed intricate relationships with the myelination status of a neuron. Specifically, Na_v_1.2 covers unmyelinated fragments in the distal axon. The myelination process itself excludes Na_v_1.2 and decreases Na_v_1.2 levels in the axon. By contrast, Na_v_1.6 only localizes to ankyrin G positive regions such as the AIS and nodes of Ranvier. In myelinated neurons, Na_v_1.6 becomes the dominant VGSC in the distal axon, as we did not detect substantial enrichment of Na_v_1.2 at nodes of Ranvier, consistent with previous results (Boiko et al., 2001; Caldwell, Schaller, Lasher, Peles, & Levinson, 2000). Na_v_1.2 and Na_v_1.6 share conserved sequence and structure homology. Thus, their abilities to establish such complex differential localization patterns are particularly intriguing.

Here, by dual labeling and 2-color super resolution imaging, we first established that, once synthesized, Na_v_1.2 and Na_v_1.6 are sorted into distinct trafficking vesicles. By coupling pulse-chase labeling with single-molecule imaging, we confirmed that the localization of Na_v_1.2 and Na_v_1.6 to AIS requires previously identified Ankyrin G-binding domain (ABD) (Garrido et al., 2003; Gasser et al., 2012; Lemaillet et al., 2003). Interestingly, we found that separated signals located in ICL1 are responsible for targeting and membrane loading of Na_v_1.2 to/at the distal axon. Strikingly, Na_v_1.6 with Na_v_1.2 ICL1 gained access to the distal axon. Na_v_1.2 ICL1 alone targets GFP molecules to cell membrane. Single molecule imaging revealed that Na_v_1.2 ICL1 promotes active transport, suppresses retention at the AIS and promotes membrane loading at the distal axon.

Our results demonstrated that the complex localization patterns of VGSCs are established by compartment-specific trafficking and loading mechanisms. For deeper understanding of molecular mechanisms, it would be critical to identify ICL1 interaction partners and their associated pathways in the future. Nonetheless, the developmental regulation and the differential localization patterns revealed in our study clarified current debates on membrane composition of VGSCs, which would help us better understand their physiological and pathological functions in the brain.

## MATERIALS AND METHODS

### Animals

Homozygous H11^LSL-Cas9^ CRISPR/Cas9 knock-in male mice (Jackson laboratory, JAX Stock #027632) (Chiou et al., 2015) were crossed with wild type C57Bl/6 females to get time pregnant heterozygous litters for *in utero* electroporation. All procedures were in accordance with protocols approved by the Janelia Research Campus Institutional Animal Care and Use Committee. Mice were housed in a 12:12 light:dark cycle.

### DNA constructs

Knockin constructs containing SpCas9, gRNA and donor DNA were modified from PX551 and PX552 backbones, which were gifts from Feng Zhang (Addgene plasmid #60957 and #60958). An EF1 promoter-driven ‘spaghetti monster’ fluorescent protein with Flag tag (smFP_Flag) (Viswanathan et al., 2015) cassette was inserted into PX552 construct to indicate successful plasmid transfection. All gRNAs were designed by CHOPCHOP (Labun et al., 2019). The gRNA targeting sequence of mouse *Scn2a* (site 1, C-terminus) is: 5’-GGACAAGGGGAAAGATATCA- 3’; The gRNA targeting sequence of rat *Scn2a* (site 2, extracellular loop between segment 5 and 6 in domain I) is: 5’-TGGTACTGCCTTCAATAGGA-3’; The gRNA targeting sequence of mouse *Scn8a* (C-terminus) is: 5’-CCGACAAGGAGAAGCAGCAG-3’. Plasmids encoding mouse Na_v_1.2 (NP_001092768.1) and Na_v_1.6 (NP_001070967.1) were cloned by Gibson Assembly (NEB) with synthetic gBlocks gene fragments (Integrated DNA Technologies). Plasmids used for electrophysiological recording tests were designed based on the final sequences after SpCas9- mediated HITI.

### Primary Culture of Hippocampal Neurons

We prepared dissociated hippocampal neurons from P0 to 1 Sprague-Dawley rat or C57Bl/6 mouse pups. Briefly, the hippocampi were dissected out and digested with papain (Worthington Biochemical). After digestion, the tissues were gently triturated and filtered with the cell strainer. The cell density was counted and ∼2.5 × 10^5^ cells were transfected with indicated constructs by using P3 Primary Cell 4D-Nucleofector X kit (Lonza). After transfection, neurons were plated onto poly-D-lysine (PDL, Sigma)-coated coverslips and maintained in NbActiv4 medium (BrainBits) at 37 °C for indicated days.

### Immunofluorescence Staining of cultured hippocampal neurons

Cultured neurons were fixed with 4% paraformaldehyde, permeabilized and blocked with 10% fetal bovine serum, 1% Triton in PBS, incubated with primary antibodies against V5 tag (R960- 25, ThermoFisher Scientific, 1:1000; 13202, Cell Signaling Technology, 1:1000), HA tag (3724, Cell Signaling Technology, 1:1000), AnkG (75-146, Antibodies Incorporated, 1:1000), MAP2 (AB5622, Millipore, 1:5000), GFP (A-11122, ThermoFisher Scientific, 1:1000), or Flag tag (ab1257, Abcam, 1:1000) overnight at 4°C. After washing with 10% fetal bovine serum in PBS, neuron samples were stained with Alexa Fluor-conjugated secondary antibodies (1:1000, ThermoFisher Scientific) and imaged with Nikon A1R confocal microscope or Zeiss LSM 880 Airyscan microscope. For actin staining, samples were stained with Alexa Fluor 594 phalloidin (A12381, 1:1000, ThermoFisher Scientific) and imaged with Leica SP8 STED microscope.

### *In Utero* Electroporation and Histology

*In utero* electroporation was performed as previously described (Mikuni et al., 2016; Petreanu, Mao, Sternson, & Svoboda, 2009). In brief, time- pregnant mouse (E13 for hippocampus and E15 for cerebral cortex) was anesthetized with 2 ∼ 2.5% isoflurane with an O_2_ flow rate of 0.5 ∼ 0.8 L/min. Before the surgery, use a cotton-tip applicator to coat both eyes with puralube and administer buprenorphine (0.1 mg/kg, intraperitoneal injection; Bedford Laboratories) for analgesia. DNA solution (1 ∼ 2 µl @ 1 µg/µl) was injected into the lateral ventricle via picospritzer. Electrical pulses (E13: 40 V for 50 ms, 8 times with 1 s intervals; E15: 45 V for 50 ms, 8 times with 1 s intervals) were delivered through ECM 830 electroporator. Administer Ketaprofen (5 mg/kg, intraperitoneal injection; Bedford Laboratories) to reduce inflammation when the surgery was done and once a day for two days after the surgery.

After mouse pups were born and reached indicated ages, they were deeply anesthetized and perfused with 4% paraformaldehyde in 0.1 M phosphate buffer, pH 7.4. The brain was dissected out and post-fixed overnight. After rinsed with PBS, coronal vibratome sections (70 µm in thickness) were made (VT1200S, Leica). The sections were permeabilized and blocked with 10% fetal bovine serum, 1% Triton in PBS, incubated with primary antibodies against V5 tag (13202, Cell Signaling Technology, 1:1000) and AnkG (75-146, Antibodies Incorporated, 1:1000), MBP (SMI-99, Millipore Sigma, 1:1000), Caspr (75-001, Antibodies Incorporated, 1:1000), CaMKIIα (MA1-048, ThermoFisher Scientific, 1:400) or GAD67 (MAB5406, Millipore Sigma, 1:1000) overnight at 4°C. After washing with 10% fetal bovine serum in PBS, neuron samples were stained with Alexa Fluor-conjugated secondary antibodies (ThermoFisher Scientific) and imaged with Zeiss 880 Airyscan microscope.

MBP staining images were used to quantify the percentage of labeled Na_v_1.2 or Na_v_1.6 in unmyelinated, myelinating and myelinated neurons. Unmyelinated neurons are the ones with Na_v_1.2 or Na_v_1.6 signals and without MBP signals along the whole axon. Partially myelinated neurons are the ones with fragmented Na_v_1.2 or Na_v_1.6 signals (> 10 µm) interspaced with MBP signals along the axon. Myelinated neurons are the ones with Na_v_1.2 or Na_v_1.6 signals in mature nodes of Ranvier (< 10 µm) interspaced with MBP signals along the axon. The intensity distribution profiles along the AIS region and the intensity levels in dendrite of Na_v_1.2 and Na_v_1.6 in mouse cortical neurons at different ages were analyzed with Fiji. The mean background intensity was subtracted before all further analysis.

### Whole-Cell Recording

HEK293 cells were cultured in Dulbecco’s modified Eagle’s culture media with 10% fetal bovine serum in a 37°C incubator with 5% CO_2_ and were grown in 60-mm culture dishes. Plasmids encoding wild type (*WT*) or V5-labeled Na_v_1.2, or wild type Na_v_1.6 or V5-labeled Na_v_1.6 (4 μg) were co-transfected with *Scn1b* (2 µg), *Scn2b* (2 µg) and eGFP (0.3 µg) using Lipofectamine 2000 (ThermoFisher Scientific). Whole-cell voltage-gated sodium (Na^+^) currents were measured 48 hours after transfection at room temperature under voltage patch-clamp configuration with an Axopatch 200B amplifier (Molecular Devices) and sampled at 10 kHz and filtered at 2 kHz. Na^+^ currents were elicited with a 50 ms depolarization step from -100 mV with 5 mV increment at a holding potential of -100 mV. Steady-state inactivation were tested by a two-pulse protocol with the first pulse of 500 ms from -100 mV to -10 mV at 5 mV increment followed by a second pulse fixed at -10 mV. Gating activation and steady-state inactivation curves were obtained using a Boltzmann function as reported previously (Wang et al., 2021). The pipette solution contained (in mM): CsF 35, CsCl 50, L-aspartic acid 55, NaCl 10, EGTA 5, MgCl_2_ 1, Mg-ATP 4, Na-GTP 0.4 and HEPES 10, pH 7.3 with CsOH; the external solution contained (in mM): NaCl 120, KCl 5.4, CaCl_2_ 1.8, MgCl_2_ 1, HEPES 10, glucose 10, tetraethylammonium chloride 20, pH 7.4 with NaOH. The access resistance was 7.9±0.9 MΩ (*WT*) versus 6.9±0.4 MΩ (V5-labeled) (*t* test, p = 0.34) with 60-80% compensation, and 6.7±0.6 MΩ (*WT*) versus 6.0±0.5 MΩ (V5-labeled) (*t* test, p = 0.43) with 80-90 % compensation for Na_v_1.2 and Na_v_1.6, respectively.

### Pulse-Chase Single Molecule Imaging

Transfected hippocampal neurons were plated onto an ultra-clean cover glass pre-coated with PDL and cultured for indicated days (days *in vitro*, DIV 9 ∼ 10). The cells were first incubated with 100 mM JF646-HTL for 1.5∼2 hrs. After washout, the labeling medium was replaced with 10 mM JF549-HTL for chase labeling (20 minutes for overexpression experiments, 40 minutes for knockin experiments). After final washout, the cover glass was transferred to live-cell culturing metal holder with phenol red free NbActiv4 medium and mounted onto Nikon Eclipse TiE Motorized Inverted microscope equipped with a 100X oil-immersion objective (Nikon, N.A. = 1.49), an automatic TIRF/HILO illuminator, a perfect focusing system, a tri-cam splitter, three EMCCDs (iXon Ultra 897, Andor) and Tokai Hit environmental control (humidity, 37 °C, 5% CO_2_). AMPPNP (1 mM, Sigma, A2647) was added during whole imaging period for indicated experiment. Before single molecule imaging, one snapshot JF646 image was captured to indicate the general labeling profile. For tracking JF549-labeled single molecules, we used 561 nm laser with the excitation power of ∼150 W/cm^2^ at an acquisition time of 100 ms.

### Fluorescent Recovery After Photobleaching

Cultured mouse hippocampal neurons harboring Na_v_1.2 or Na_v_1.6 knockin with HaloTag (DIV 9 ∼ 10) were labeled with 20 mM JF549-HTL for 0.5 hr. After washout, the cover glass was transferred to live-cell culturing metal holder with phenol red free NbActiv4 medium and mounted onto Zeiss LSM 880 confocal microscope equipped with a 40X oil-immersion objective (N.A. = 1.40), a definite focus module, and a large incubation unit for global CO_2_ and temperature control (37 °C) along with heated stage insert. 3 frames were acquired before photobleaching and 297 frames were acquired to observe fluorescent recovery after photobleaching with a time interval of 15 seconds. Pitstop 2 (30 µM, Sigma, SML1169) was added to inhibit endocytosis during the labeling and imaging period for indicated conditions. Fiji was used to analyze fluorescent recovery. Relative intensity of the photobleached region of interest (ROI) were calculated by subtracting mean intensity of the background from the photobleached ROI region, followed by normalized to the mean intensity of the pre-bleach ROI.

### Single-Molecule Localization, Tracking and Diffusion Analysis

For single-molecule localization and tracking, the spot localization (x,y) was obtained through 2D Gaussian fitting based on MTT algorithms (Serge, Bertaux, Rigneault, & Marguet, 2008). The localization and tracking parameters in SPT experiments are listed in the **Table 1**. The Radius of Confinement (RC) for each trajectory is calculated as the distance between the center of mass (the average position of all localizations in the trajectory) to the furthest localization from the center of mass. The differential probability density function (PDF) curve is obtained by subtraction of RC PDF distributions between conditions as indicated in each figure panel.

**Table 1.**
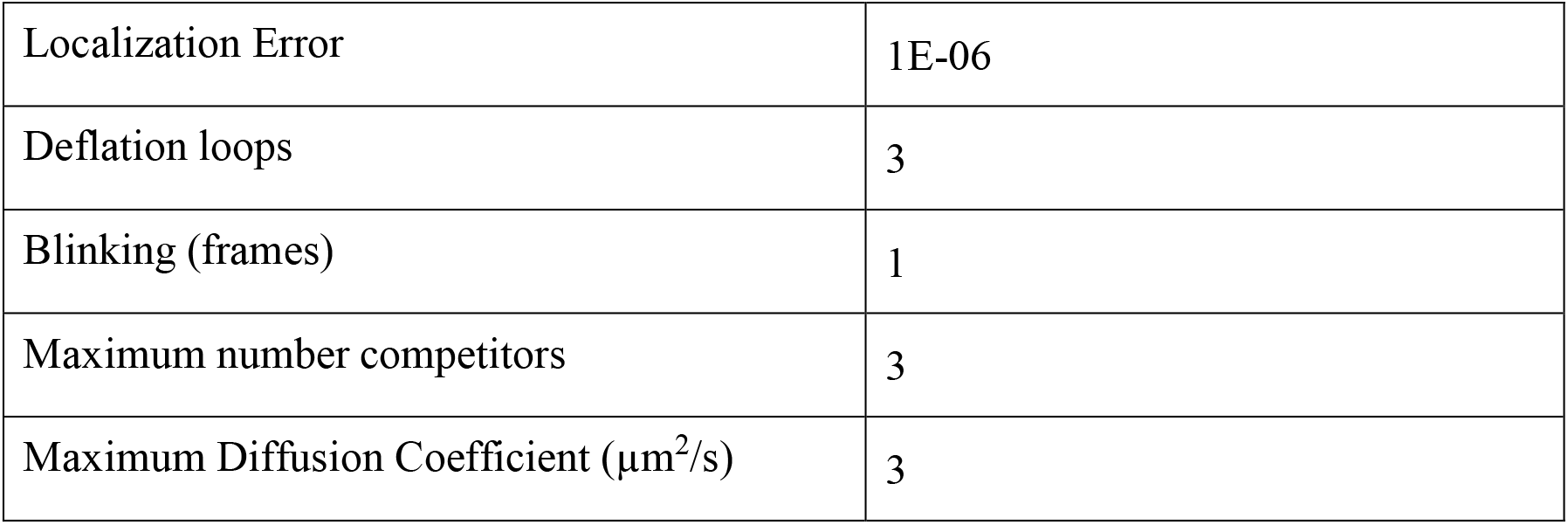
Localization and tracking parameters for the MTT program

### Statistics

Comparisons between two groups were performed with Student’s *t* test. Comparisons among multiple groups were performed with one-way ANOVA and *post hoc* Bonferroni test. Differences were considered to reach statistical significance when p < 0.05.

### Data Availability Statement

The data that support the findings of this study are available from the corresponding author upon request.

## AUTHOR CONTRIBUTIONS

Z.J.L. and H.L. conceived and designed the experiments. H.L. performed and participated in all experiments. G.S.P. and H.G.W. designed and performed the electrophysiology experiment. Z.J.L. and H.L. analyzed the data and wrote the manuscript. G.S.P. and H.G.W. helped write the manuscript. Z.J.L. supervised the research.

---

**Figure 1-figure supplement 1.**
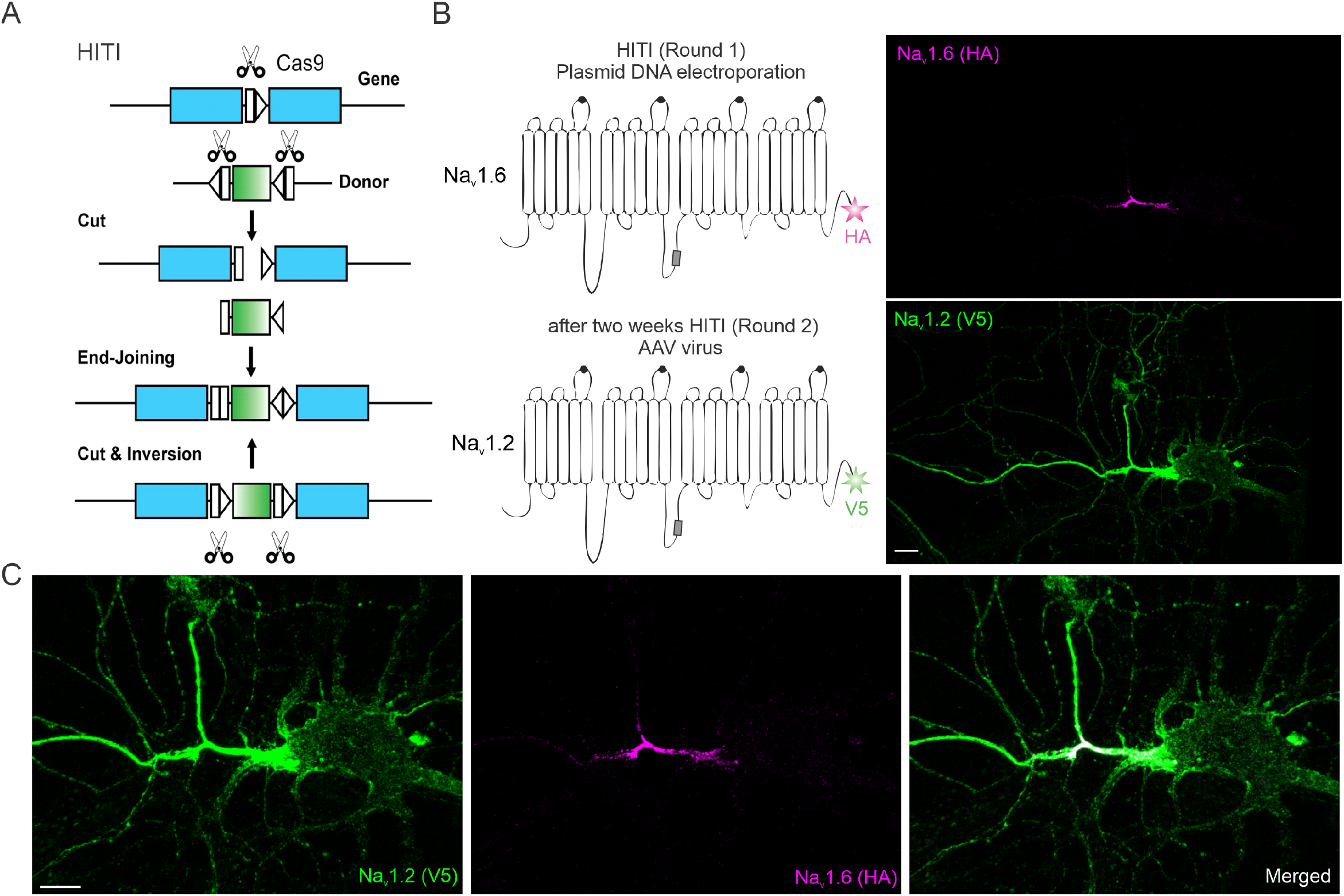
The HITI knock-in strategy and dual labeling of Na_v_1.2 and Na_v_1.6 in the same neuron. (**A**) The schematics for the HITI strategy. The donor DNA fragment has two gRNA cutting sites flanking the tag cDNA. After cutting and end-joining, if the fragment is inserted into the genome in the right direction, the two gRNA cutting sites will be inactivated. If not, the cutting and end-joining process will continue until it is inserted in the right direction. (**B**) The strategy of double- labeling Na_v_1.2 and Na_v_1.6 in the same neuron (left). Through sequential plasmid DNA electroporation and AAV virus infection two weeks later, double-labeling of Na_v_1.2 and Na_v_1.6 in the same neuron is achieved. The neuron in Figure 1A is shown in a larger field of view on the right. (**C**) Zoom-in views of the soma, dendrite and AIS region of the example neuron. Scale bar in (B) and (C), 10 µm.

**Figure 1-figure supplement 2.**
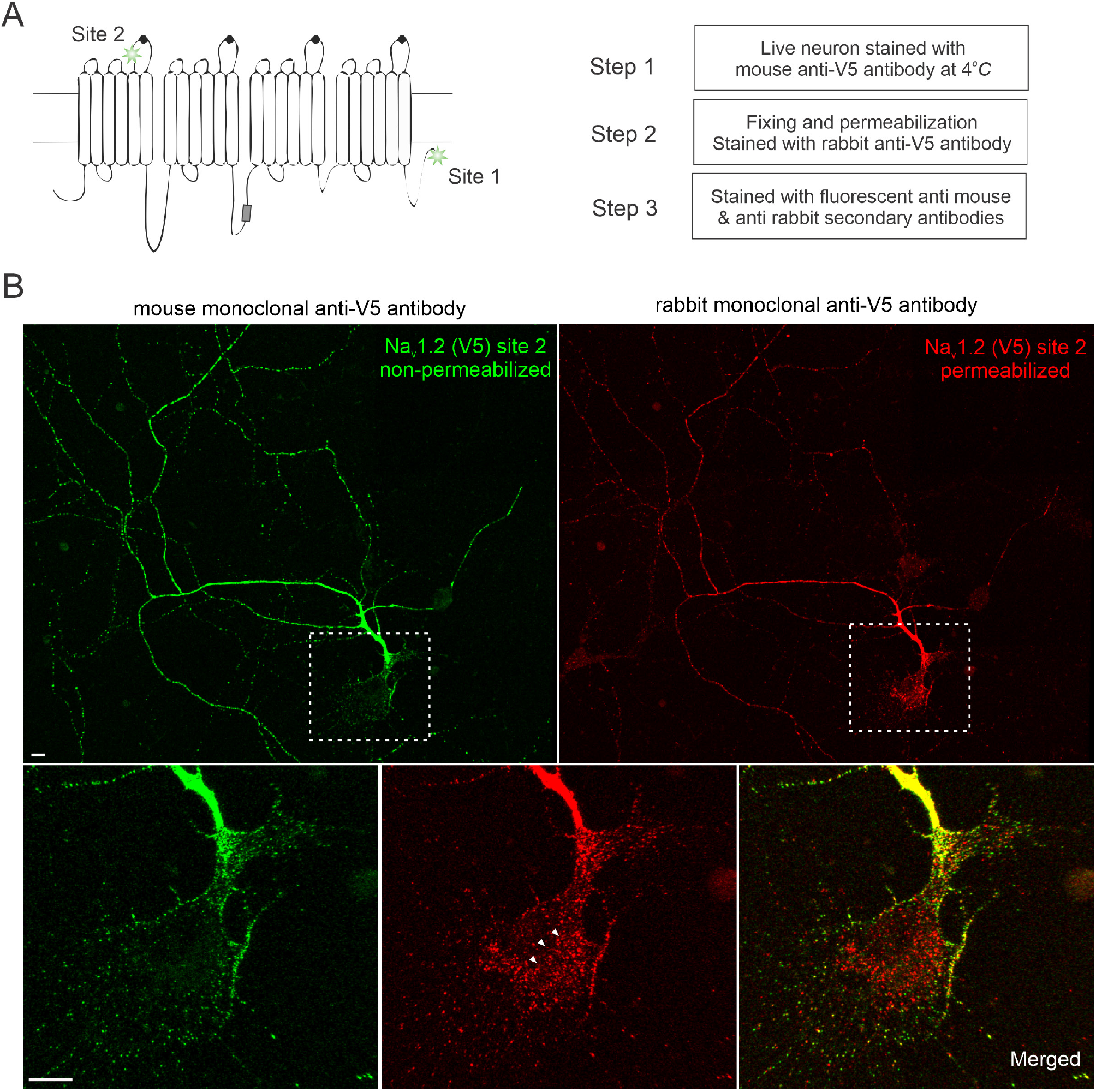
Non-permeabilized staining of Nav1.2 confirms its membrane localization at the distal axon and dendrites. (**A**) Left shows two gRNA targeting sites of Na_v_1.2 used in this study. Site 2 is at the extracellular loop between segment 5 and 6 of domain I. With V5 insertion at this site, we performed three-step staining shown in the right protocol. (**B**) Non-permeabilized (green) and permeabilized staining (red) images of Na_v_1.2 in the same neuron. Bottom images are zoom-in views of the soma region (rectangle region with dashed lines). Arrowheads indicate labeled intracellular Na_v_1.2 vesicles (red) which were absent in the non-permeabilized staining (green). Scale bar, 10 µm.

**Figure 1-figure supplement 3.**
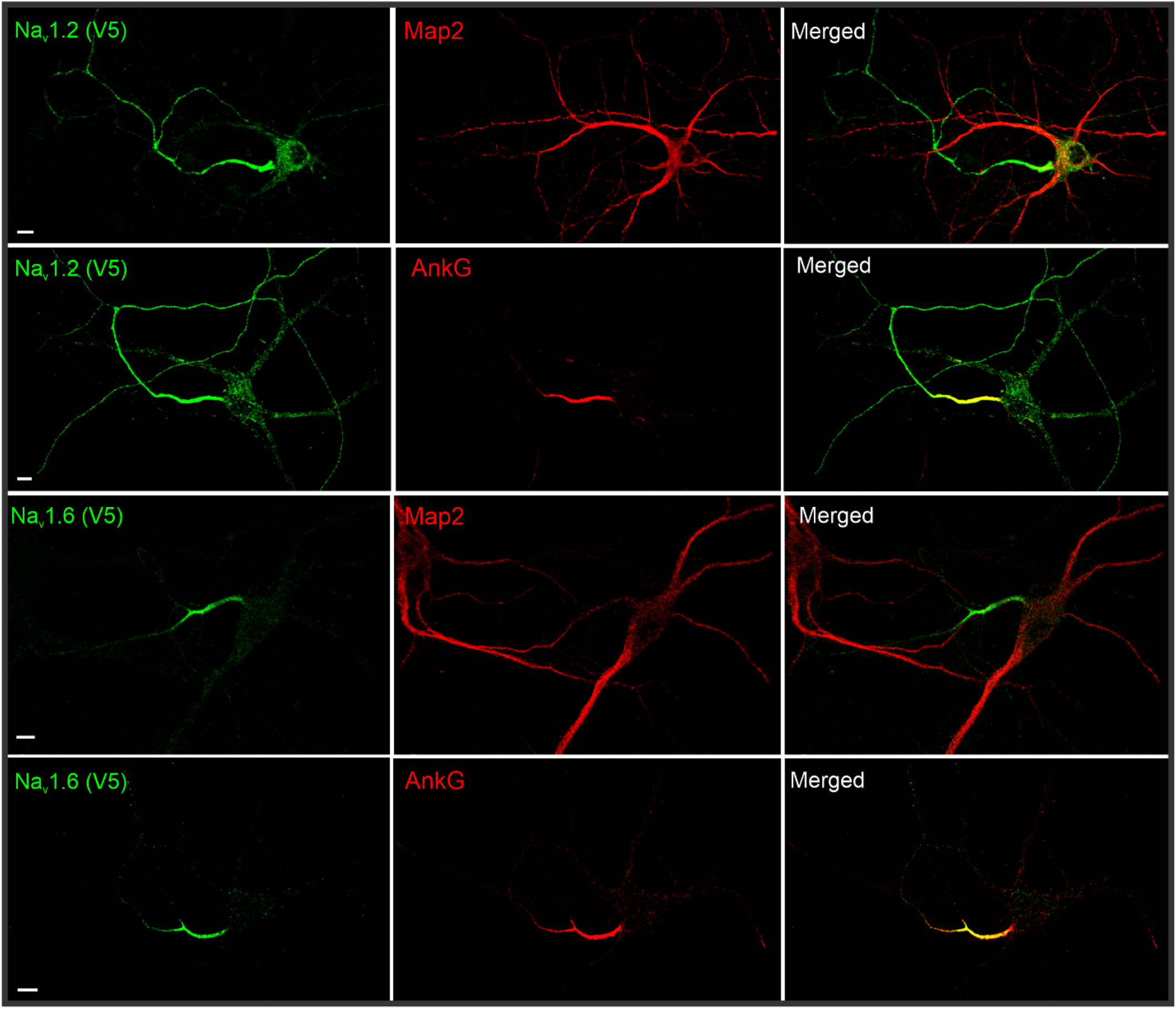
Immunostaining of V5-labeled Na_v_1.2 and Na_v_1.6 with MAP2 or AnkG in cultured hippocampal neurons. By cross referencing with dendrite (MAP2) and AIS (AnkG) markers, we found that Na_v_1.2 is enriched in AIS, distal axon and dendrites, whereas Na_v_1.6 is mainly localized in the AIS region. Scale bar, 10 µm.

**Figure 1-figure supplement 4.**
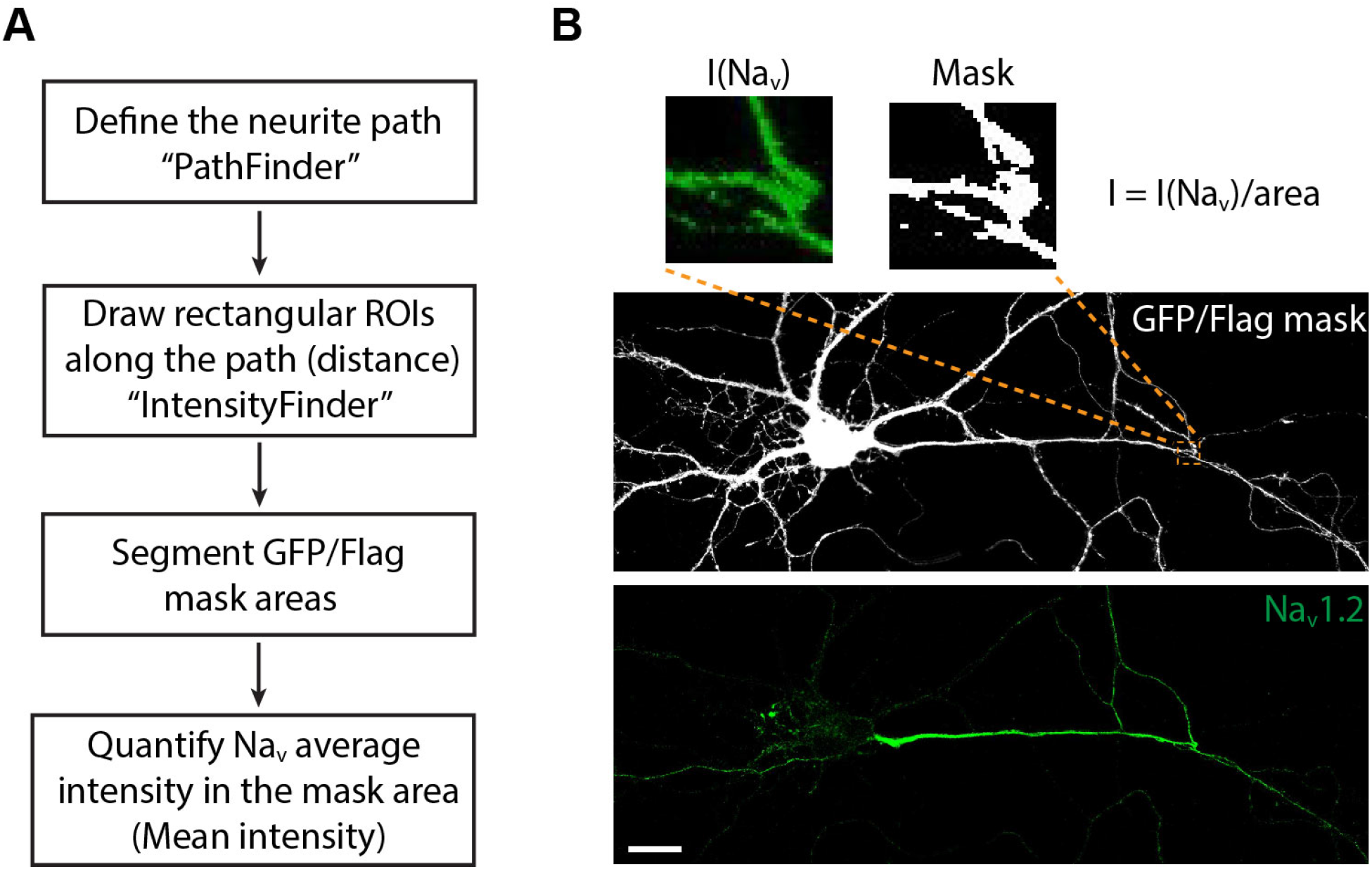
An analysis pipeline for estimating Na_v_1.2 and Na_v_1.6 sub- cellular abundance. (**A**) First, the analysis pipeline defines the neurite path (axon and dendrite); secondly, ROIs were selected along the path with the calculation of their distances from the soma; lastly, the program segments GFP/Flag mask areas and quantifies Na_v_ average intensities in the neurites within each ROI (Figure 1-figure supplement 4-source code 1, 2). (**B**) An example showing segmentation and quantification in one ROI. The mean intensity is calculated through normalization of total Na_v_ intensity with the mask area. Scale bar, 20 µm.

**Figure 1-figure supplement 5.**
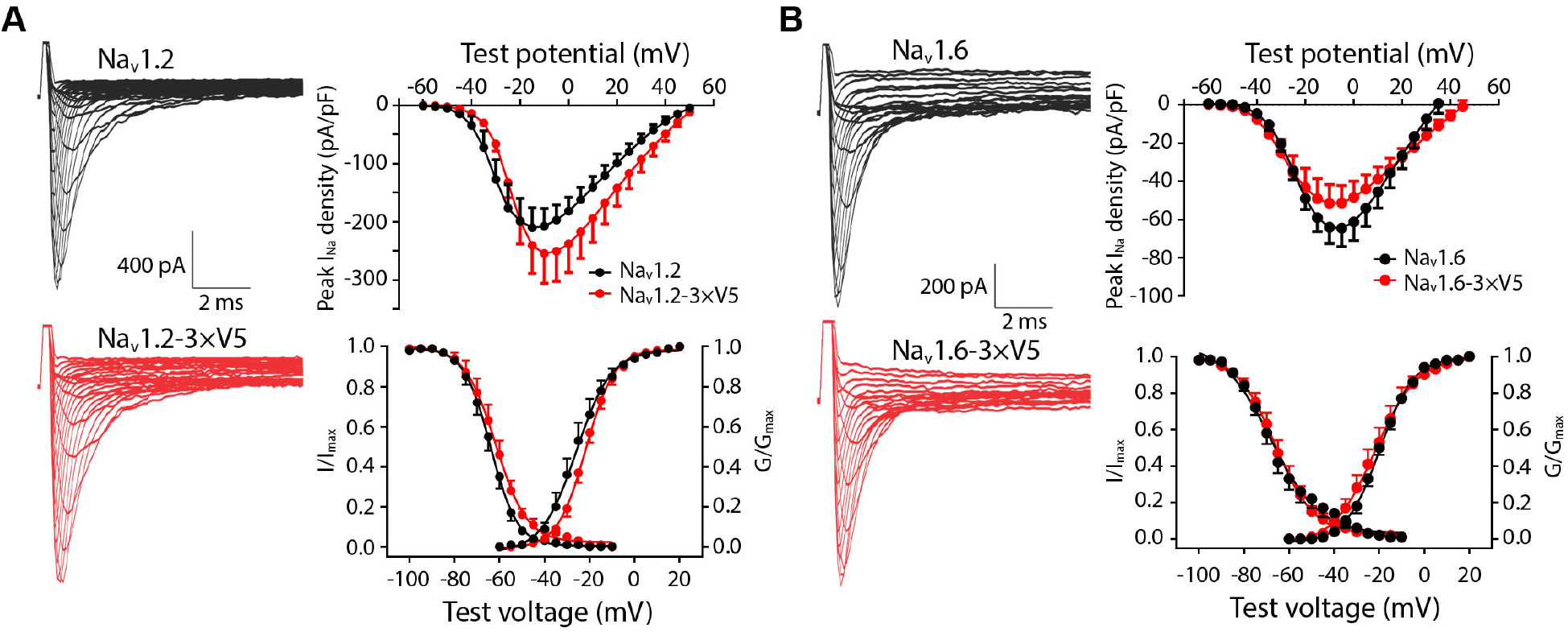
Electrophysiological properties of wild type and V5-labeled Na_v_1.2 and Na_v_1.6 in HEK293T cells. (**A**) *WT* (black) and V5-labeled (red) Na_v_1.2. Left: Na_v_1.2 current examples; right: peak current density (upper), channel activation (*WT*, n = 10; V5-labeled, n = 9) and steady-state inactivation (*WT*, n=11; V5-labeled, n = 9) curves (lower). (**B**) *WT* (black) and V5-labeled (red) Na_v_1.6. Left: Na_v_1.6 current examples; right: peak current density (upper), channel activation (*WT*, n = 11; V5- labeled, n = 12) and steady-state inactivation (*WT*, n = 13; V5-labeled, n = 11) curves (lower). See in Figure 1-figure supplement 5-source data 1.

**Figure 2-figure supplement 1.**
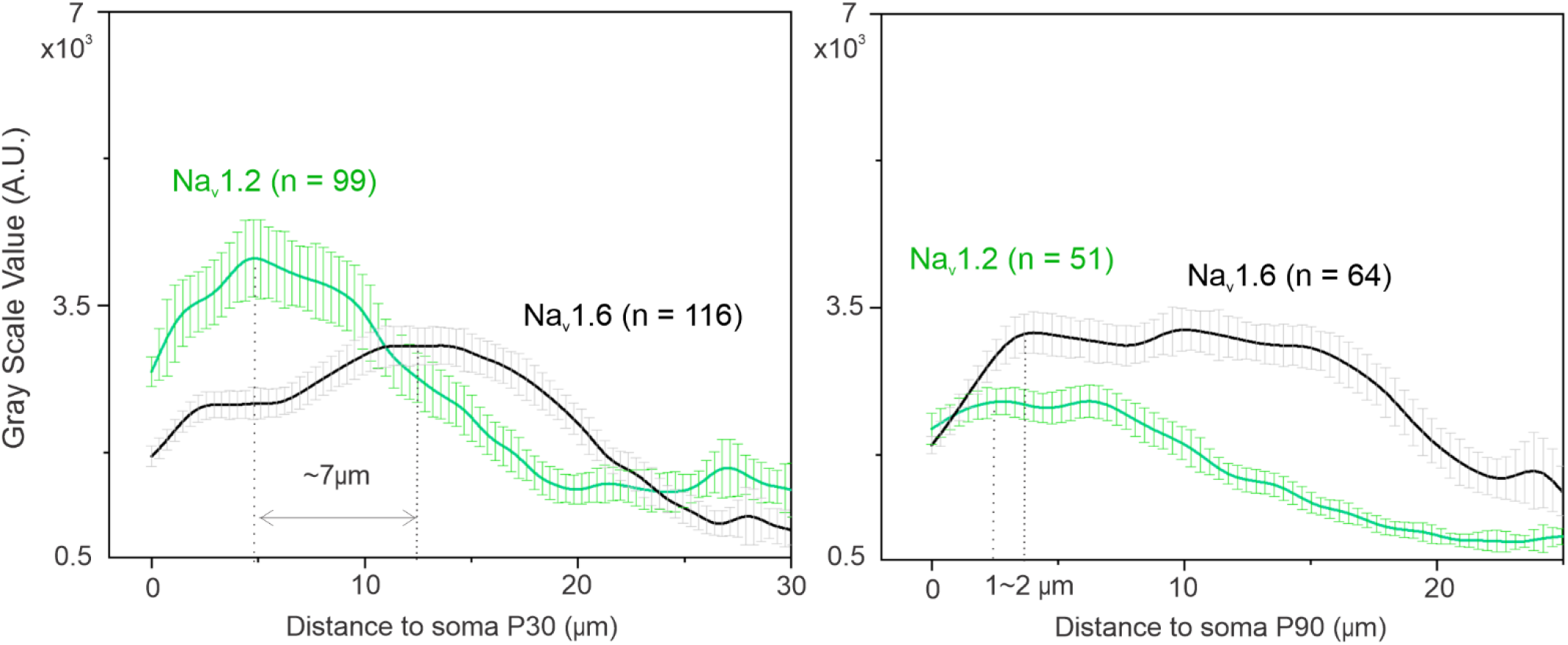
The distribution profiles of Na_v_1.2 and Na_v_1.6 along the AIS of mouse cortical neurons at P30 and P90. From P15 (Figure 2F) to P90, Na_v_1.2 levels in AIS gradually decrease and the concentration peak of Na_v_1.6 shift inwards, moving closer to the Na_v_1.2 concentration peak located at the proximal AIS. See in Figure 2-figure supplement 1-source data 1, 2. Error bars represent SEM; n indicates the number of cells analyzed.

**Figure 2-figure supplement 2.**
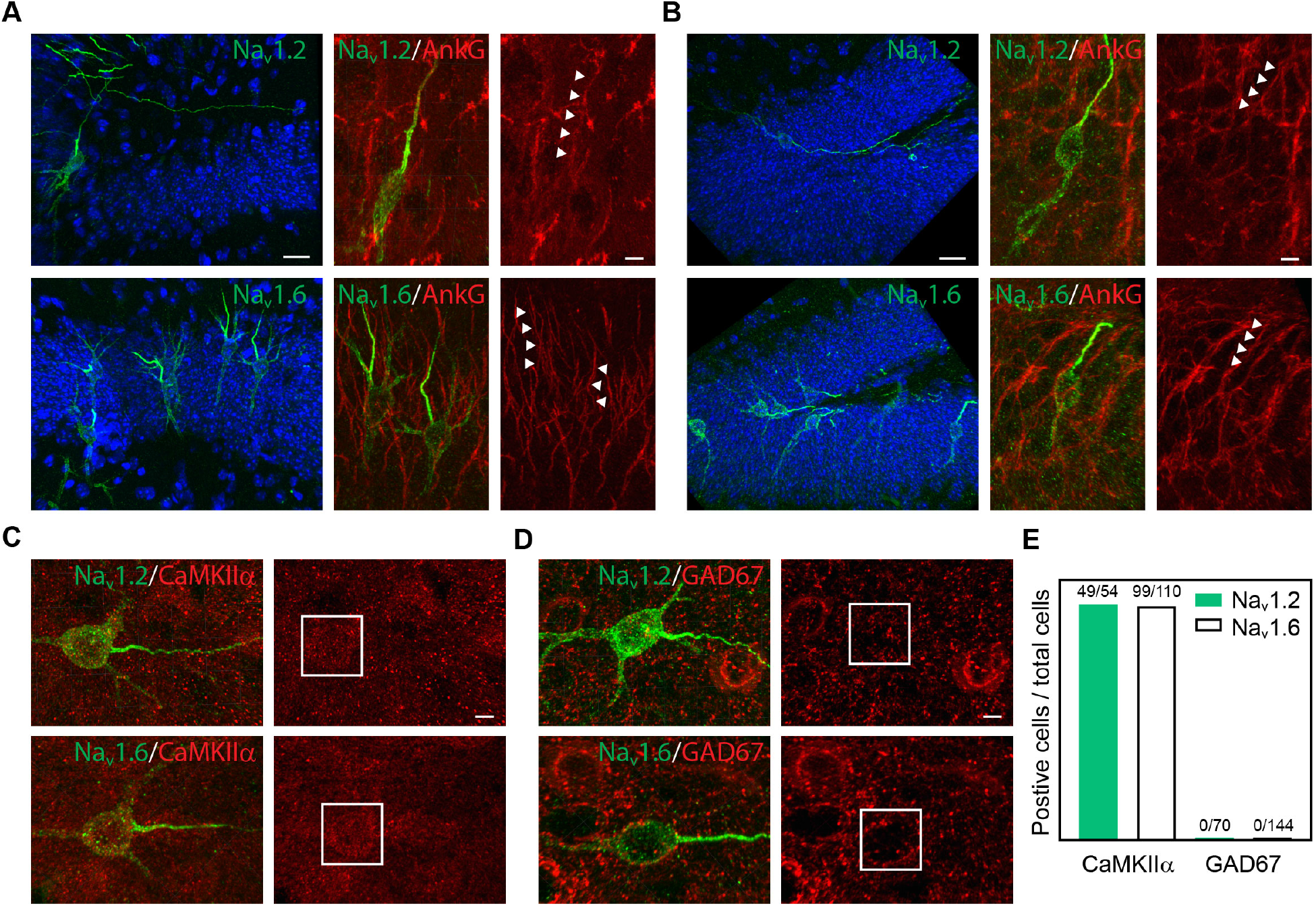
Cell-type specific expression of Na_v_1.2 and Na_v_1.6 in the mouse brain. (**A**) and (**B**) Representative images of Na_v_1.2 and Na_v_1.6 labeled with V5 tag in CA1 (A) and dentate gyrus (B) of the hippocampus. Left: Blue channel shows the Hoechst stain. Scale bar, 20 µm. Right: Zoom-in images co-stained with AnkG. Arrowheads indicate Na_v_1.2 or Na_v_1.6-positive region with AnkG signals. Scale bar, 5 µm. (**C**) and (**D**) Double-immunostaining of V5-labeled Na_v_1.2 or Na_v_1.6 with CaMKIIα (C) or GAD67 (D) in the mouse cortex. Rectangles highlight the soma regions of Na_v_1.2 or Na_v_1.6-positive neurons. Scale bar, 5 µm. (**E**) The positive ratio of Na_v_1.2 or Na_v_1.6 knockin cells in CaMKIIα-positive excitatory or GAD67-positive inhibitory neurons.

**Figure 3-figure supplement 1.**
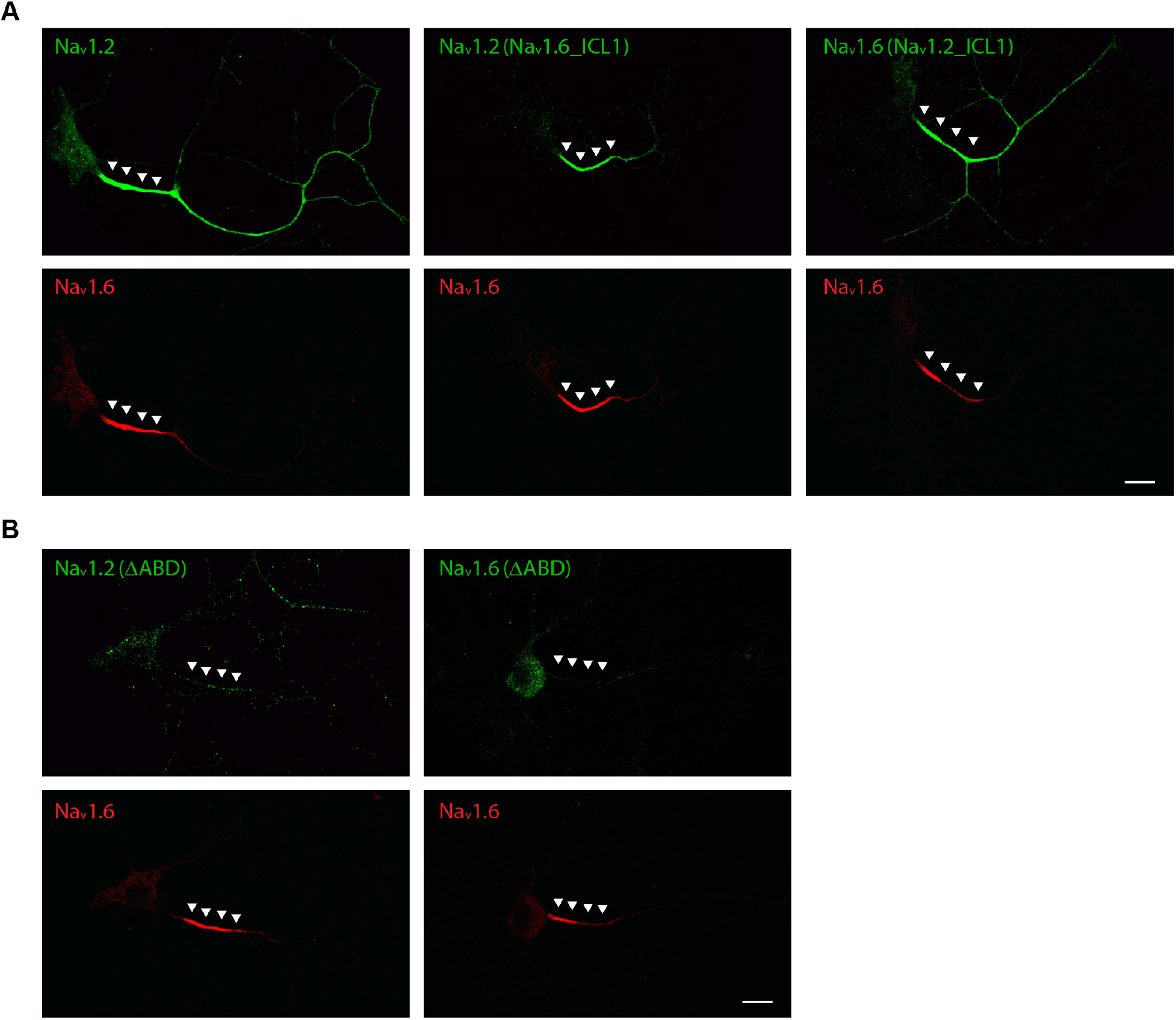
Localization of Na_v_1.2 to the distal axon requires ICL1. (**A**) Left: localization patterns of wild-type Na_v_1.2 (green) and Na_v_1.6 (red). Middle: Na_v_1.2 with Na_v_1.6 ICL1 showed minimal enrichment at the distal axon. Right: Na_v_1.6 with Na_v_1.2 ICL1 gained access to the distal axon. (**B**) ABD deletion greatly reduced Na_v_1.2 and Na_v_1.6 levels at the AIS. Arrowheads indicate the AIS region. Scale bar, 20 µm.

**Figure 3-figure supplement 2.**
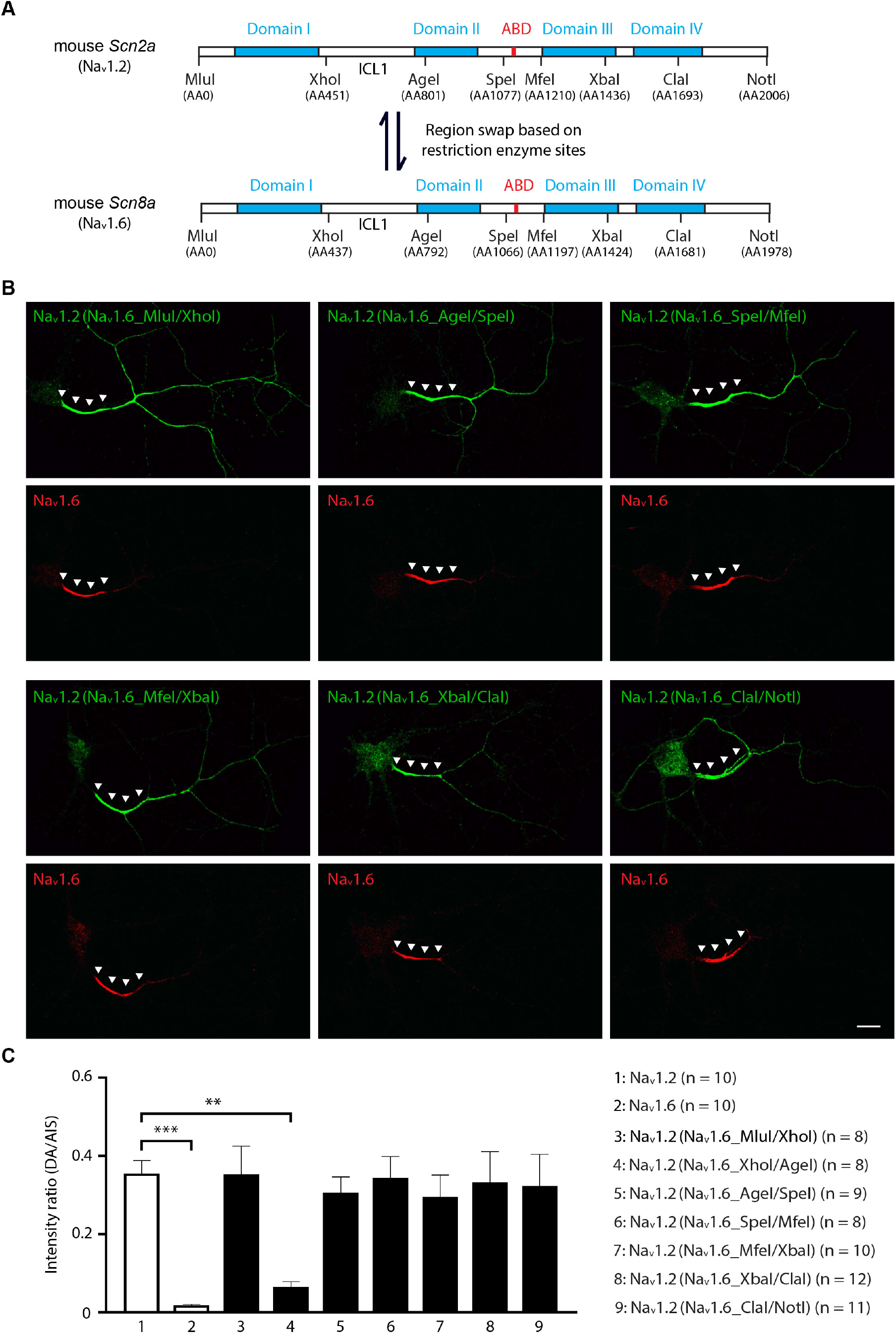
Identification of the domain required for the localization of Na_v_1.2 to the distal axon. (**A**) Eight restriction enzyme sites were designed in both mouse *Scn2a* (Na_v_1.2) and *Scn8a* (Na_v_1.6) cDNA without altering their protein sequences. Subcloning was used to swap corresponding regions between Na_v_1.2 and Na_v_1.6. ICL1 is indicated in the graph, which could be exchanged with XhoI and AgeI. (**B**) Swapping the regions other than ICL1 domain (XhoI and AgeI) did not affect Na_v_1.2 localization in the distal axon (green). ICL1 representative images are shown in Figure 3**-figure supplement 1A**. Arrowheads indicate the AIS region. Scale bar, 20 µm. (**C**) Analysis of the intensity ratio between distal axon and AIS of Na_v_1.2 with swapped domains from Na_v_1.6 (Figure 3-figure supplement 2-source data 1). Error bars represent SEM. ***, *p*-value < 0.001; **, *p*-value < 0.01.

**Figure 3-figure supplement 3.**
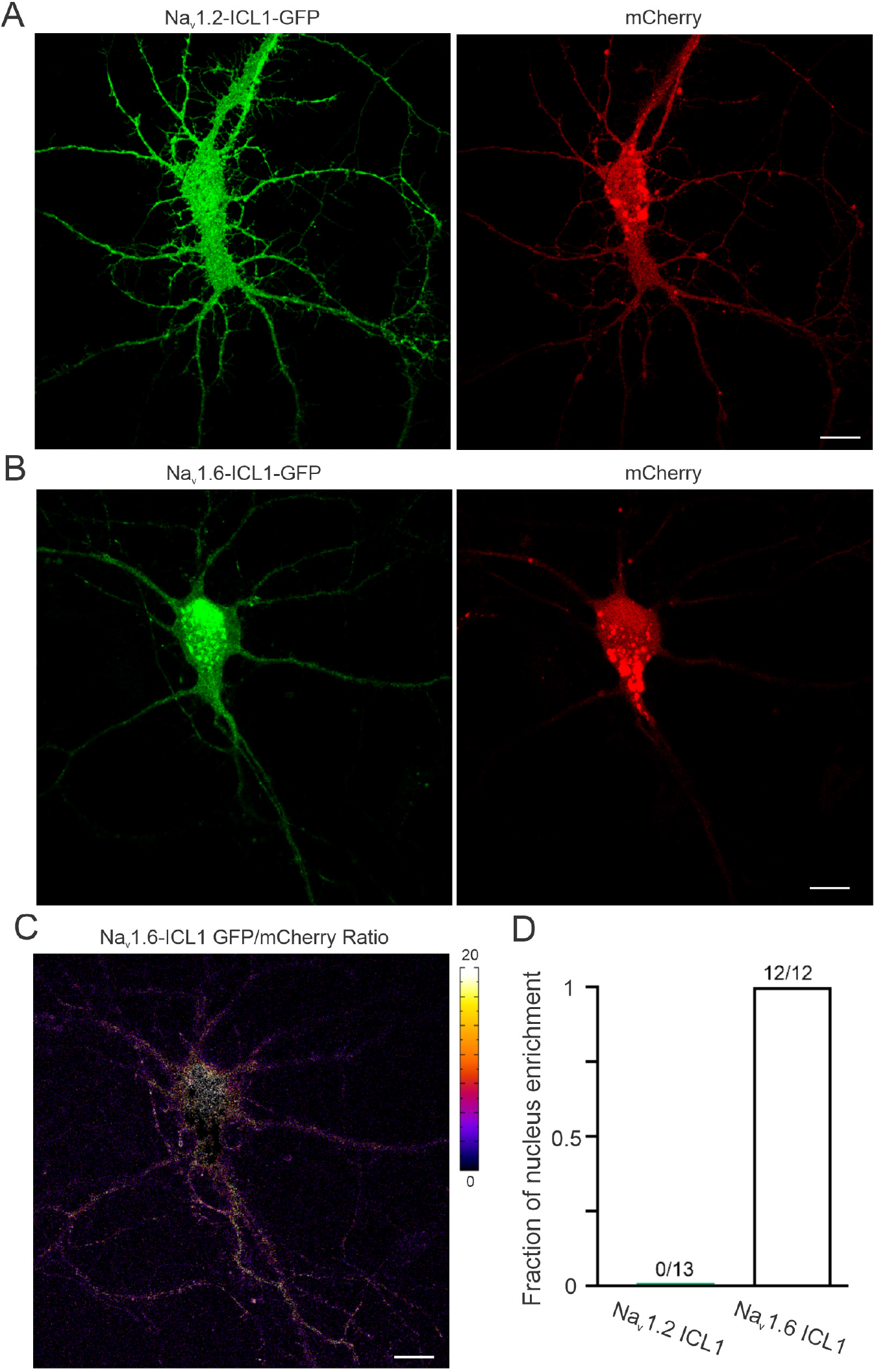
Na_v_1.2 ICL1 and Na_v_1.6 ICL1 target GFP to membrane and the nucleus respectively. (A) and (**B**) Representative raw images of Na_v_1.2 (A) and Na_v_1.6 (B) ICL1-GFP and corresponding mCherry signals. (**C**) Intensity ratio image of Na_v_1.6 ICL1-GFP/mCherry shown in (B) suggests a nucleus enrichment of Na_v_1.6-ICL1-GFP. Right color bar indicates the ratio level. (**D**) The percentage of neurons showed nucleus localization of GFP signals. Scale bar (A - C), 20 µm.

**Figure 3-figure supplement 4.**
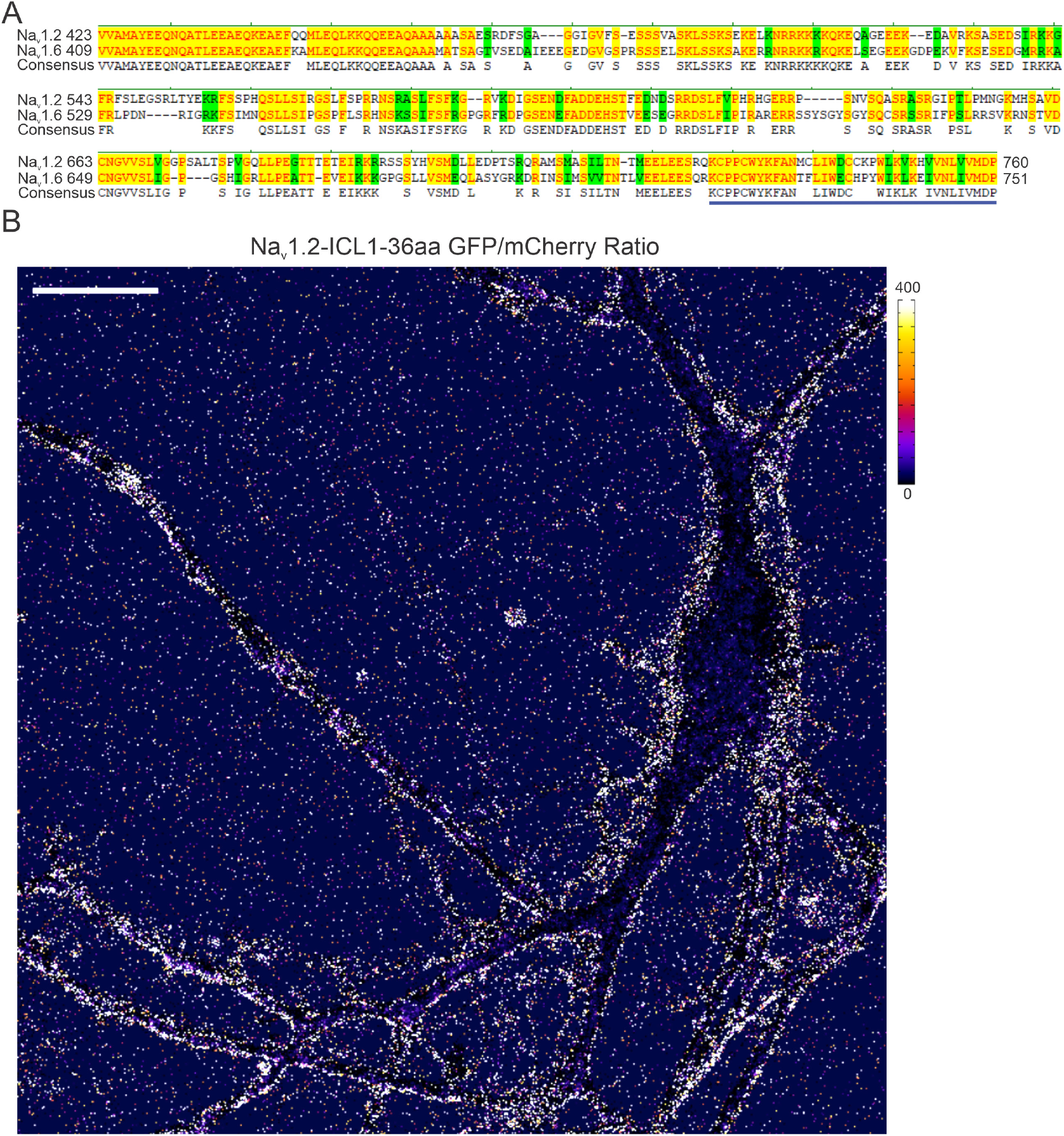
Na_v_1.2 ICL1-36aa targets GFP to membrane. (**A**) Protein sequence alignment of Na_v_1.2 and Na_v_1.6 ICL1 region. Bold blue line highlights identified 36aa region. (**B**) A representative image of Na_v_1.2 ICL1-36aa GFP/mCherry ratio showed the specific distribution of Na_v_1.2 ICL1-36aa along the membrane. Right color bar indicates the ratio level. Scale bar, 20 µm.

**Figure 4-figure supplement 1.**
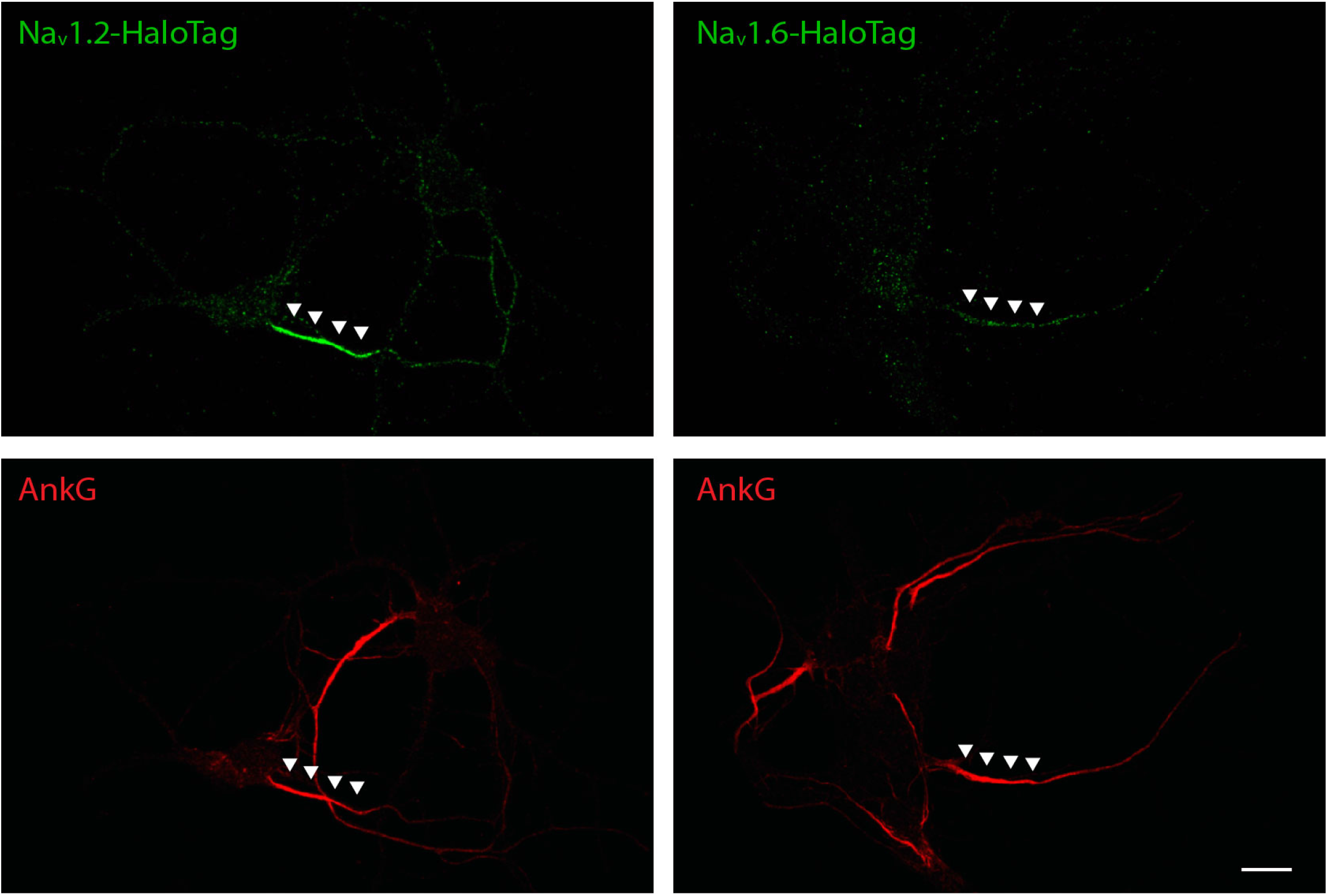
HaloTag-labeled Na_v_1.2 and Na_v_1.6 were enriched in AIS. With JF646-HTL bulk labeling, Na_v_1.2 (left) and Na_v_1.6 (right) with HaloTag knockin have higher concentrations in the AIS, similar to V5 or HA tag knockin. Scale bar, 10 µm.

## VIDEOS

**Video 1 (Related to Figure 1)**

A representative cultured hippocampal neuron with double labeling of Na_v_1.2 (V5, green) and Na_v_1.6 (HA, red) co-stained with Flag (blue).

**Video 2 (Related to Figure 1)**

Raw Airyscan image of V5-labeled Na_v_1.2 (green) and HA-labeled Na_v_1.6 (purple) vesicles in the soma region of a cultured hippocampal neuron and their distributions after segmentation. Scale bar, 5 µm.

**Video 3 (Related to Figure 2)**

3D Airyscan image of Na_v_1.2 knockin neurons (V5, green) in mouse cortex co-stained with MBP (red) shows its continuous distribution along distal axons without myelin coverage.

**Video 4 (Related to Figure 2)**

3D Airyscan image of Na_v_1.6 knockin neurons (V5, green) in mouse cortex co-stained with MBP (red) shows its presence in myelinated neurons.

**Video 5 (Related to Figure 2)**

3D Airyscan image of Na_v_1.6 knockin neurons (V5, green) in mouse cortex co-stained with Caspr (red) shows its localization at nodes of Ranvier.

**Video 6 (Related to Figure 4)**

FRAP experiment of a cultured hippocampal neuron with HaloTag-labeled Na_v_1.2 (stained with JF549-HTL, red) shows very slow fluorescent recovery at AIS (white rectangle) after photobleaching and moderate fluorescent recovery in distal axons (green and yellow rectangles). Scale bar, 10 µm.

## SOURCE DATA AND CODE FILES

**Figure 1-source data 1 (Related to Figure 1D)**

**Figure 1-source data 2 (Related to Figure 1E)**

**Figure 1-figure supplement 4-source code 1, 2 (Related to Figure 1-figure supplement 4A)**

**Figure 1-figure supplement 5-source data 1 (Related to Figure 1-figure supplement 5)**

**Figure 2-source data 1 (Related to Figure 2F)**

**Figure 2-source data 2 (Related to Figure 2G)**

**Figure 2-figure supplement 1-source data 1, 2 (Related to Figure 2-figure supplement 1)**

**Figure 3-source data 1 (Related to Figure 3B)**

**Figure 3-source data 2 (Related to Figure 3C)**

**Figure 3-figure supplement 2-source data 1 (Related to Figure 3-figure supplement 2C)**

**Figure 4-source data 1 (Related to Figure 4C)**

**Figure 4-source data 2 (Related to Figure 4D)**

**Figure 4-source data 3 (Related to Figure 4E)**

**Figure 4-source data 4 (Related to Figure 4F)**

## ACKNOWLEDGEMENTS

We thank M. Radcliff for assistance and Liu Lab members for providing help and suggestions. We thank L. Sarah, the breeding and surgery team at Janelia Research Campus for helping us with *in utero* electroporation experiment. We thank W. Deepika for helping us with primary culture of hippocampal neurons. We thank the histology facility and advanced imaging center at Janelia Research Campus for helping us with histology and imaging experiments. We thank Lavis Lab at Janelia Research Campus for kindly providing JF549-HTL and JF646-HTL. Z.J.L. and H.L. is funded by Howard Hughes Medical Institute (HHMI). G.S.P. and H.G.W. is funded by NIH R01 MH118934.

